# Identification of novel broad host-range promoter sequences functional in diverse *Pseudomonadota* by a promoter-trap approach

**DOI:** 10.1101/2024.06.26.600856

**Authors:** Diego M. Roldán, Vanesa Amarelle

## Abstract

Finding novel promoter sequences is a cornerstone of synthetic biology. To contribute to the expanding catalog of biological parts, we employed a promoter-trap approach to identify novel sequences within an Antarctic microbial community that act as broad host-range promoters functional in diverse *Pseudomonadota*. Using *Pseudomonas putida* KT2440 as host, we generated a library comprising approximately 2,000 clones resulting in the identification of thirteen functional promoter sequences, thereby expanding the genetic toolkit available for this chassis. Some of the discovered promoter sequences prove to be broad host-range as they drove gene expression not only in *P. putida* KT2440 but also in *Escherichia coli* DH5α, *Cupriavidus taiwanensis* R1^T^, *Paraburkholderia phymatum* STM 815^T^, *Ensifer meliloti* 1021, and an indigenous Antarctic bacterium, *Pseudomonas* sp. UYIF39. Our findings enrich the existing catalog of biological parts, offering a repertoire of broad host-range promoter sequences that exhibit functionality across diverse members of the phylum *Pseudomonadota,* proving Antarctic microbial community as a valuable resource for prospecting new biological parts for synthetic biology.

## 1. Introduction

Promoters are key elements in gene expression and regulation. They are an invaluable genetic tool in synthetic biology (SynBio) and genetic engineering, as they are fundamental building blocks for the assembly of synthetic biological circuits and for recombinant protein expression, among other uses. Most promoter sequences used in SynBio were designed based on *Escherichia coli* promoters. This is reflected by the available promoter sequences in the Registry of Standard Biological Parts, where more than three hundred promoters are from *E. coli*, in comparison with the seven promoters available from *Bacillus subtilis*, and only nine promoters from other prokaryotes ^1^.

While *E. coli* remains the most used chassis in SynBio, the tendency is to evaluate and implement non-model bacteria as alternatives, to overcome *E. coli* limitations ^3^. In this context, the availability of broad host-range promoters is of interest to favour compatibility among different chassis. Despite the significance of such promoters, limited attention has been given to identifying/constructing broad-host range promoters and assessing their functionality in different bacterial host. Some of the few examples in this field include the work by Schuster *et al.* who generated a toolbox intended for *Pseudomonadota* that includes twelve promoter–regulator pairs ^4^, most of them originally from *E. coli*. The toolkit was evaluated in nine strains of the phylum, representing α, β and γ *Proteobacteria,* reporting some broad-host range inducible promoters ^5^. Yang *et al.* developed three constitutive promoters of different strengths, functional in *E. coli*, *Bacillus subtilis*, and *Saccharomyces cerevisiae* ^6^. In a recent work, Keating and Young evaluated standard parts in different *Pseudomonadota*, obtaining different promoter activity depending on the host assessed ^7^. All these examples evaluated canonical promoter sequences, rationally designed based on *E. coli* promoter architecture, mainly those sequences involved in σ^70^ binding. With the advent of non-model chassis, finding novel sequences in nature that exert promoter activity in a broad-range of species is of great interest. In this context, function-driven metagenomics has a lot to offer. Mainly used to search for new enzymatic activities ^8^, this approach can also be applied for identification of regulatory elements. Despite its potential, only few works used this approach, either to find novel promoter sequences functional in *E. coli* ^9–11^, or transcriptional terminators functional in *Pseudomonas putida* and other *Pseudomonadota*^12^.

In the last few years, *P. putida* became a reliable SynBio chassis for designing and building genetic circuits for a plethora of applications ^13^. There are different tools available for this chassis, including promoter sequences ^14^. Most constitutive promoters used in *P. putida* are originally from *E. coli*, with only few works where promoter sequences specifically intended for this chassis were developed ^15,16^. In this context, finding new promoter sequences functional in *P. putida* is of great interest.

The aim of this study was to identify and characterise new versatile promoters functional in different *Pseudomonadota* using a function-driven metagenomic approach. Instead of using *E. coli* as host, we used *P. putida* KT2440 to find novel promoter sequences for this SynBio chassis, killing two birds with one stone. By means of a promoter-trap vector with *gfp* as reporter and Antarctic soil metagenomic DNA as input, we identified thirteen novel promoter sequences functional not only in *P. putida* KT2440, but also in other members of the phylum *Pseudomonadota*. This work expands the SynBio toolbox by identifying and characterizing novel broad host-range promoters.

## 2. Materials and methods

### 2.1. Bacterial strains, plasmids, and growth conditions

Bacterial strains and plasmids used in this study are detailed in Table S1. All strains, with the exception of *Ensifer meliloti* 1021, were grown aerobically at 25 °C and 200 rpm in LB medium (10 g/L tryptone, 5 g/L yeast extract, 10 g/L NaCl), or in M9 minimal medium (47.7 mM Na_2_HPO_4_, 22 mM KH_2_PO_4_, 8.6 mM NaCl_2_, 18.7 mM NH_4_Cl, 2 mM MgSO_4_, 0.1 mM CaCl_2_) supplemented with 0.2 % (w/v) casamino acid and 0.4% (w/v) glucose as the sole carbon source. When required, 50 μg/mL kanamycin (Km) was added to LB (LBKm) or M9 (M9Km) medium. *E. meliloti* 1021 was grown aerobically at 25 °C and 200 rpm in TY medium (5 g/L tryptone, 3 g/L yeast extract, 0.5 mM CaCl_2_) or in M9 minimal medium supplemented with 0.4% (w/v) glucose, 6 mM glutamate, 200 μM methionine and 1 μM biotin (M9S). When required, 100 μg/mL streptomycin (Str) and/or 50 μg/mL neomycin (Nm) was added to TY (TYStr, TYNm, TYStrNm) or M9S (M9SStr, M9SNm, M9SStrNm) medium.

### 2.2. Promoter-trap plasmid

In a previous work, we generated plasmid pSEVA231-*gfp* containing a promoterless *gfplva* gene ^17^ which we used in this work as a promoter-trap. Briefly, the construction is based on pSEVA231 as scaffold, therefore harbouring the pBBR1 origin of replication and Km^R^ as selectable marker. In pSEVA231-*gpf* plasmid, the *gfplva* gene has a ribosome binding site (RBS) and a BamHI recognition site 9 bp and 54 bp upstream of its start codon, respectively. Here, the BamHI site was used for cloning the metagenomic DNA fragments in order to identify and retrieve sequences capable of exerting expression of the *gfplva* gene.

### 2.3. Antarctic soil sampling and metagenomic DNA extraction

Soil samples were collected at the Fildes Peninsula of King George Island, South Shetland archipelago, Antarctica. Sampling was performed during the February 2020 campaign and soil samples were kept at −20 °C in sterile tubes until processing. The samples were collected in the following sites: 62°10’13.8’’S, 58°55’42.6’’W; 62°10’06.6’’S, 58°56’06.4’’W; and 62°10’16.3’’S, 58°55’35.0’’W. Metagenomic DNA from each soil sample was extracted using the Quick-Start Protocol DNeasy^Ⓡ^ PowerSoil^Ⓡ^ Kit (QIAGEN), as suggested by the manufacturer. DNA integrity was determined by agarose gel electrophoresis, and their quantity and quality were determined spectrophotometrically.

### 2.4. Promoter-trap metagenomic library construction

Metagenomic DNA of the three soil samples were pooled, and partial digestion with Sau3AI was optimized in order to obtain fragments of 0.1-0.5 kb. Briefly, starting with 6.0 µg of metagenomic DNA, 5 and 10 U of enzyme, and 5, 10, 15, 30 and 45 min digestion times were assessed. The digestion reaction was stopped by thermal inactivation at 65 °C for 20 min. The efficiency of the procedure was evaluated by agarose gel electrophoresis. When fragments from appropriate size (0.1-0.5 kb) were obtained, they were excised and extracted from the agarose gel with QIAquick Gel Extraction Kit (QIAGEN). The metagenomic fragments were cloned into the dephosphorylated BamHI site of pSEVA231-*gpf*. Ligation mixtures were used to transform *P. putida* KT2440 cells as previously reported ^17^. Briefly, cells were grown in LB until late exponential phase, were washed thrice with sterile distilled water at room temperature, and were finally resuspended in 1:20 of the original volume. One hundred microliters of competent cells were electroporated with 1 µL of the ligation mixture and then 900 µL of LB were added. Cells were recovered for 1h at 25 °C and 100 µL were plated in LBKm. Plates were incubated for 16 h at 25 °C, and clones expressing the green fluorescent protein (GFP) were evidenced using a Safe Imager^TM^ 2.0 blue-light transilluminator (Invitrogen). Positive clones were selected and re-streak in LBKm plates. Plasmids were isolated and were used to re-transform *P. putida* KT2440 cells in order to confirm the observed phenotype.

### 2.5. Introduction of constructs in members of the phylum Pseudomonadota

In order to evaluate if the metagenomic DNA sequences that proved to act as functional promoter sequences in *P. putida* KT2440 were also functional in other members of the phylum *Pseudomonadota*, the thirteen constructs (generically called pSEVA231-*Pr(x)gfp*) were introduced into *Pseudomonas* sp. UYIF39, *Cupriavidus taiwanensis* R1^T^, *Paraburkholderia phymatum* STM 815^T^, *Escherichia coli* DH5α and *Ensifer meliloti* 1021. In the case of *Pseudomonas* sp. UYIF39, *C. taiwanensis* R1^T^, and *P. phymatum* STM 815^T^, electrocompetent cells were obtained as mentioned before and the constructs were introduced by electroporation. In the case of *E. coli* DH5α, chemical competent cells were obtained as standard protocols ^18^. Constructs pSEVA231-*Pr(x)gfp* were introduced in the different host either by electroporation of heat shock, being used in all cases 100 µl of competent cells and 10 ng of plasmid DNA. Transformants were plated in LBKm, incubated at 25 °C, and GFP expression was qualitatively determined as mentioned before.

To introduce the constructs in *E. meliloti* 1021, a triparental maiting approach was used. *E. coli* DH5α harbouring the different constructs (DH5α pSEVA231-*Pr(x)gfp*) were used as donor strains and *E. coli* DH5α harbouring plasmid pRK2013 was used as helper strain ^19^. Briefly, all strains were grown until late exponential growth phase in either LBKm at 37 °C (for donor and helper strains) or TYStr at 30 °C (for recipient strain). Cultures were diluted 1:100 in either LB or TY (as appropriate) without antibiotic, and were grown until early exponential phase (OD_600nm_=0.3). Matings were performed by mixing 100 µL of each strain, centrifuging the mix for 5 min at 5,000 rpm, and suspending it in 20 µL of TY broth without antibiotics. The mix was placed as a drop on TY agar plates and incubated at 30 °C for 24 h. Cells were collected with a loop, suspended in 1 mL of TY broth, and 100 µL were plated in TYStrNm. Transconjugants of *E. meliloti* 1021 pSEVA231-*Pr(x)gfp* were evaluated for GFP expression qualitatively as mentioned before. Appropriate mating controls were performed.

### 2.6 In vivo characterization of promoter activity

Promoter activity was determined in the different hosts at 25 °C. Despite this temperature is not the optimal for many of the strains, it is a temperature where all of them grow properly and therefore is suitable for comparison among strains. For this purpose, strains harbouring plasmids pSEVA231-*Pr(x)gfp* were grown in LBKm medium at 25 °C for 16 h (or 48 h in TYStrNm for *E. meliloti* 1021) and cultures were diluted 1:20 (v/v) in M9Km medium (or M9StrNm for *E. meliloti* 1021) in 96-well microplates, in a final volume of 200 μL. Strains were grown at 25 °C for 60 h, when optical density (OD_600nm_) and promoter activity (GFP) were determined. To quantify GFP expression, a 450 nm excitation filter and 505 nm emission filter were used. Promoter relative activity is expressed as GFP emission at 505 nm normalized to optical density (reported as GFP_505nm_/OD_600nm_). Parental strains and strains harbouring the pSEVA231-*gfp* plasmid were used as negative controls. In pSEVA231-*gfp*, the *gfp* gene lacks a promoter sequence and therefore is not being expressed. This plasmid provides a baseline for basal fluorescence that might occur even in the absence of a dedicated promoter. The parental strains establish the baseline for putative intrinsic fluorescence of the strains. Strains harbouring plasmids pSEVA231-*Pj100gfp*, pSEVA231-*Pj106gfp* and pSEVA231-*Pj114gfp* were used as controls to determine promoter activity strength. In these plasmids, *gfp* expression is under the control of Anderson promoters J23100 (here *Pj100*, strong strength promoter), J23106 (here *Pj106,* medium strength promoter), and J23114 (here *Pj114,* low strength promoter) ^20^. These canonical promoters allow us to evaluate the relative strength of the retrieved promoters in terms of GFP expression ^17^. All strains and constructs were assessed in triplicates and the experiment was repeated at least two times. Differences in GFP expression were evaluated by analysis of variance (ANOVA). The significances of the mean differences were estimated with the Tukey test (*p* < 0.05).

### 2.7 Sequencing and in silico analysis

Amplification of promoter sequences cloned in plasmids pSEVA231-*Pr(x)gfp* was performed using primers PS1 5’-AGGGCGGCGGATTTGTCC-3’ and PS2 5’-GCGGCAACCGAGCGTTC-3’ from pSEVA231 scaffold, which flank the BamHI site were the metagenomic DNA fragments were cloned. Amplicons were sequenced in both strands and were manually assembled. Sequences were used for taxonomic affiliation using PhylopythiaS Web Server ^21^. Putative open reading frames (ORFs) were identified using the online ORF Finder server in NCBI website (https://www.ncbi.nlm.nih.gov/orffinder/). Nucleotide and amino acid sequences were compared against the NCBI public database using the basic local alignment search tool (BLAST) at the NCBI on-line server.

For promoter prediction, three different platforms were used: i) Bacterial promoter prediction (BacPP) ^22^; ii) Sequence Analyser for the Prediction of Prokaryote Homology Inferred Regulatory Elements (SAPPHIRE) ^23^; and iii) Neural Network Promoter Prediction (BDGP) of Berkeley *Drosophila* Genome Project ^24^.

To determine conserved motifs present in the promoter sequences, the WebLogo platform (http://weblogo.berkeley.edu/logo.cgi) was used in promoter sequences predicted with the different algorithms (BacPP, SAPPHIRE, BDGP). For comparison, we also identified conserved motifs in the promoter regions of *Pseudomonas* spp. by retrieving all RpoD-dependent promoter sequences available in SAPPHIRE database.

The similarity between the σ^70^ factor of *P. putida* KT2440 with the σ^70^ factors of the other strains was evaluated. The assemblies were annotated using Rapid Annotation Subsystem Technology (RAST 2.0) ^25^, and the similarity between predicted σ70 coding sequences of the different strains and the σ70 protein of *P. putida* KT2440 was determined. The genomes of *P. putida* KT2440 (GCF_900167985.1), *Pseudomonas* sp. UYIF39 (GCF_029963765.1), *E. coli* DH5α (GCF_019444045.1), *C. taiwanensis* R1^T^ (GCF_000069785.1), *P. phymatum* STM 815^T^ (GCF_902833665.1), and *E. meliloti* 1021 (GCF_000006965.1) were used in this approach.

### 2.8 Selected sequences as proof of concept

In order to determine which promoter is leading *gfp* expression in sequences where multiple putative promoter sequences were identified, we selected and synthesize sequences of 50 bp based on the *in silico* results and cloned them upstream of *gfp*. In order to achieve the double strand sequence, forward and reverse synthetic primers (Table S2) were annealed by a temperature gradient (5 min at 95 °C, 72 cycles with a 1 °C decrease per cycle, during 1 min each). Primers were designed with EcoRI/BamHI sites in order to clone the annealed fragments in pSEVA231-*gfp.* Selected promoter sequences were also used to elucidate if the different promoter strength (in terms of GFP expression) exerted by the different metagenomics fragments is related to the distances of the putative promoter sequences regarding *gfp*.

Constructs harboring the selected sequences were introduced in *P. putida* KT2440 and promoter strength was qualitatively and quantitatively determined as mentioned before. In all cases, correct cloning was verified by sequencing.

## 3. Results

### 3.1 Thirteen promoter sequences functional in P. putida KT2440 were retrieved from Antarctic metagenomic DNA

Using Antarctic soil metagenomic DNA sequences as input and GFP protein as reporter, a promoter-trap metagenomic library of 1,804 clones was generated in *P. putida* KT2440. By this approach, thirteen promoters leading *gfp* expression were identified (Fig. 1).

**Figure 1.**
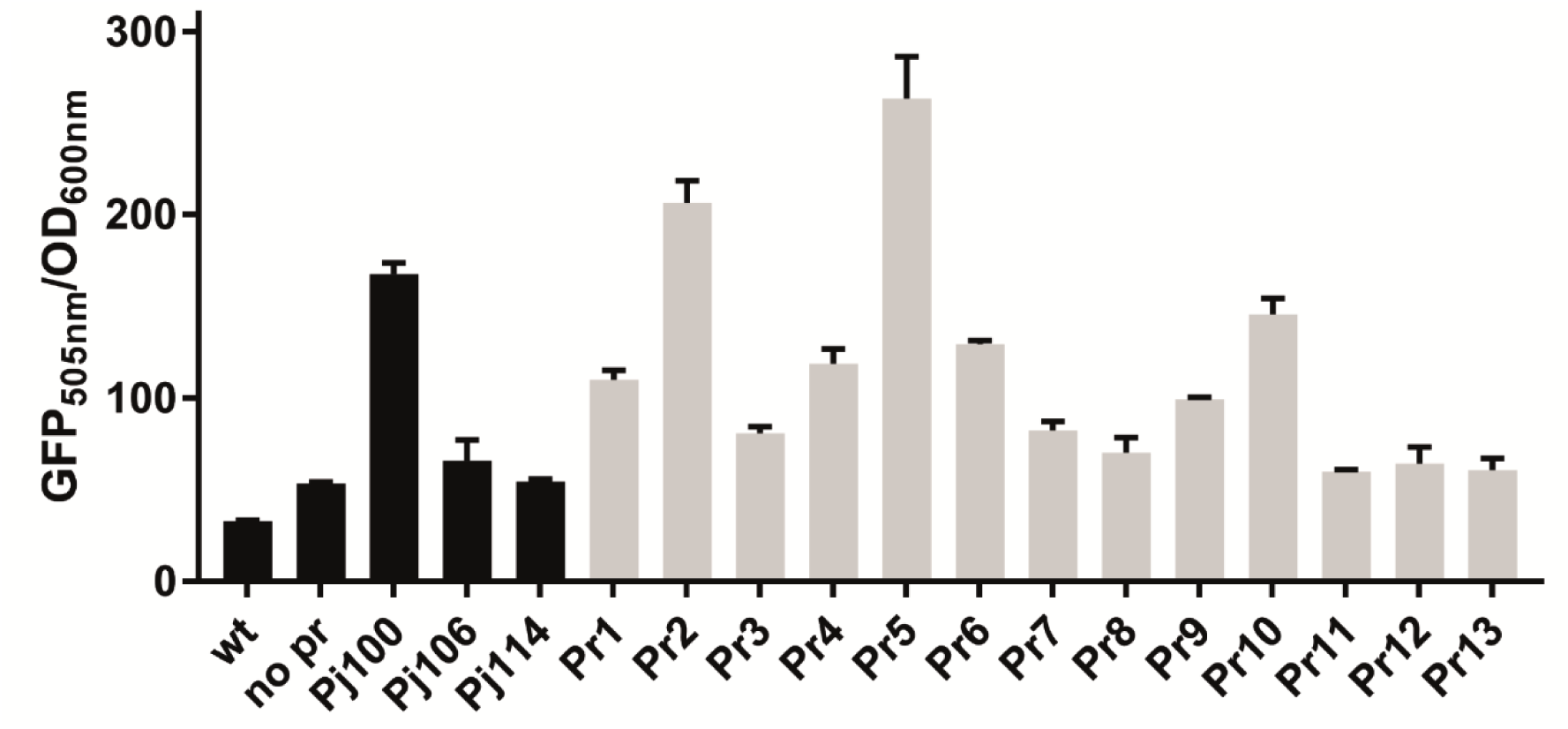
Functional promoter sequences in *P. putida* KT2440 retrieved from Antarctic metagenomic DNA. The functionality of the new promoters (*Pr1-Pr13*) was evaluated with respect to the parental strain *P. putida* KT2440 (wt) and *P. putida* KT2440 harbouring promoter-trap plasmid without promoter (no pr) as negative controls. *P. putida* KT2440 harbouring plasmids with Anderson promoters *Pj100*, *Pj106*, and *Pj114* were used as controls to determine promoter strength. Quantitative GFP expression was determined in M9Km broth, at 25 °C, and at 60 h growth. Fluorescence was normalized to OD_600nm_ for each strain. Results are the mean of three technical replicates in the same experiment. The experiment was repeated at least two times with similar results. Differences in GFP expression were evaluated by analysis of variance (ANOVA). The significances of the mean differences were estimated with the Tukey test (*p* < 0.05).

As shown in Fig. 1, sequences *Pr2* and *Pr5* exerted an expression that exceeds that of the canonical strong promoter *Pj100,* and were therefore categorized as strong promoters. Sequences *Pr1*, *Pr4*, *Pr6*, *Pr9* and *Pr10* exerted an expression in-between *Pj100* and *Pj106* canonical promoters. Sequences *Pr3*, *Pr7*, *Pr8*, exerted an expression similar to *Pj106* and were considered medium strength promoters, while sequences *Pr11*, *Pr12* and *Pr13* expression barely exceeded that of promoter *Pj114* and are therefore considered weak promoters. None of the constructs generated a noteworthy metabolic burden to *P. putida* KT2440 as evidenced by the similar profiles of growth curves (Fig. S1).

### 3.2 In silico analysis of the metagenomic inserts

To gain insight into the promoter sequences retrieved by the promoter-trap approach, all 13 constructs were sequenced and analyzed *in silico*. Sequences were deposited in GenBank with accession numbers OQ288876-OQ288888.

As shown in Table 2, the average insert size is 465 bp, being the longest sequence *Pr8* (968 bp) and the shortest sequence *Pr7* (242 bp). Considering the insert size average, the whole library (1,804 clones) can be estimated to represent 838 kb, which is equivalent to approximately 0.18 genomes considering an average genome size of 4.5 Mb ^26^. Taxonomic affiliation of the inserts by PhylopythiaS determined that they mainly belong to the phyla *Actinomycetota* (6 sequences) and *Pseudomonadota* (5 sequences), and the remaining sequences belong to the phyla *Acidobacteriota* (*Pr10*) and ‘*Candidatus* Methylomirabilota’ (*Pr13*). Inserts show a mean G+C content of 60.4%, in a range of 56-66%.

**Table 2.**
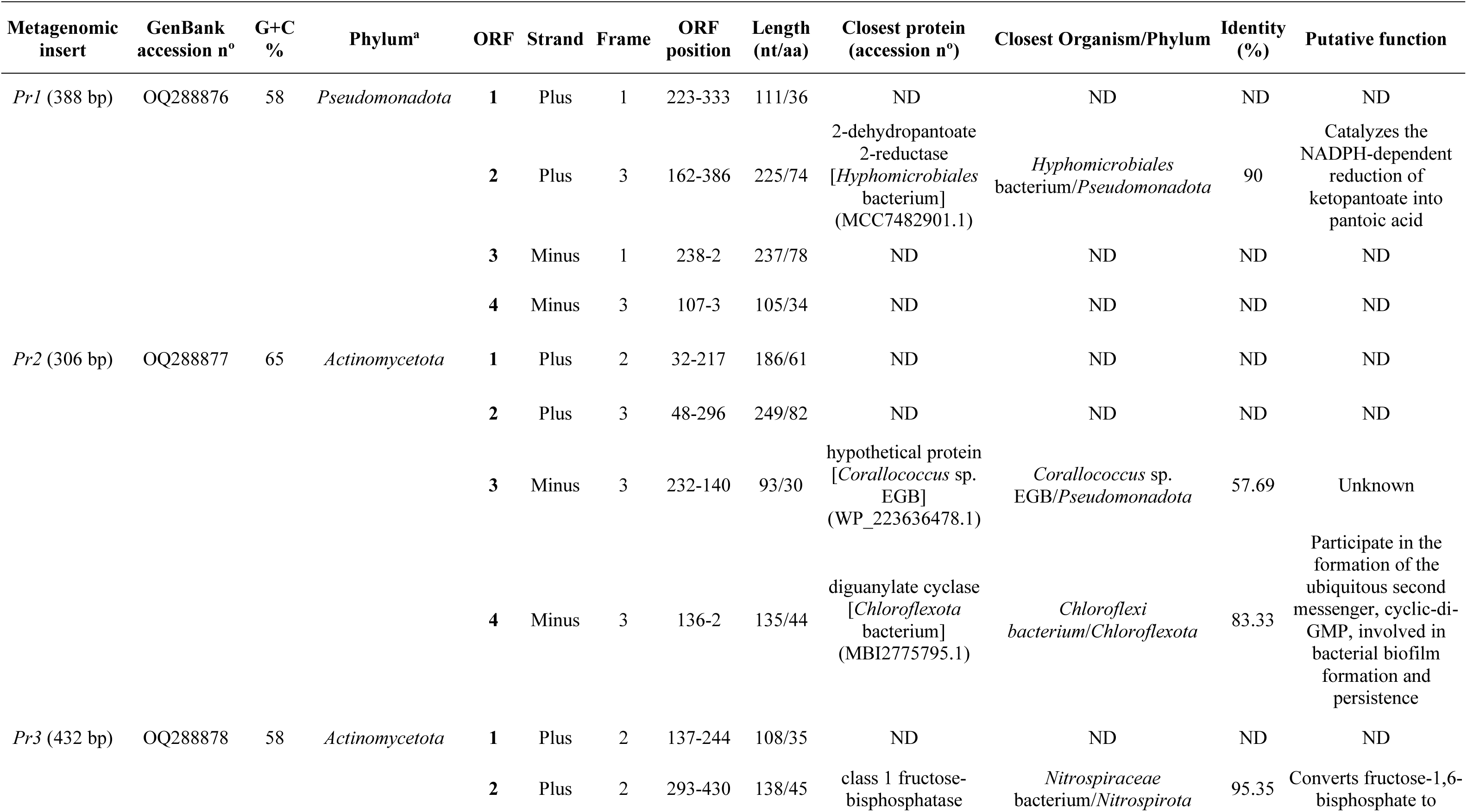
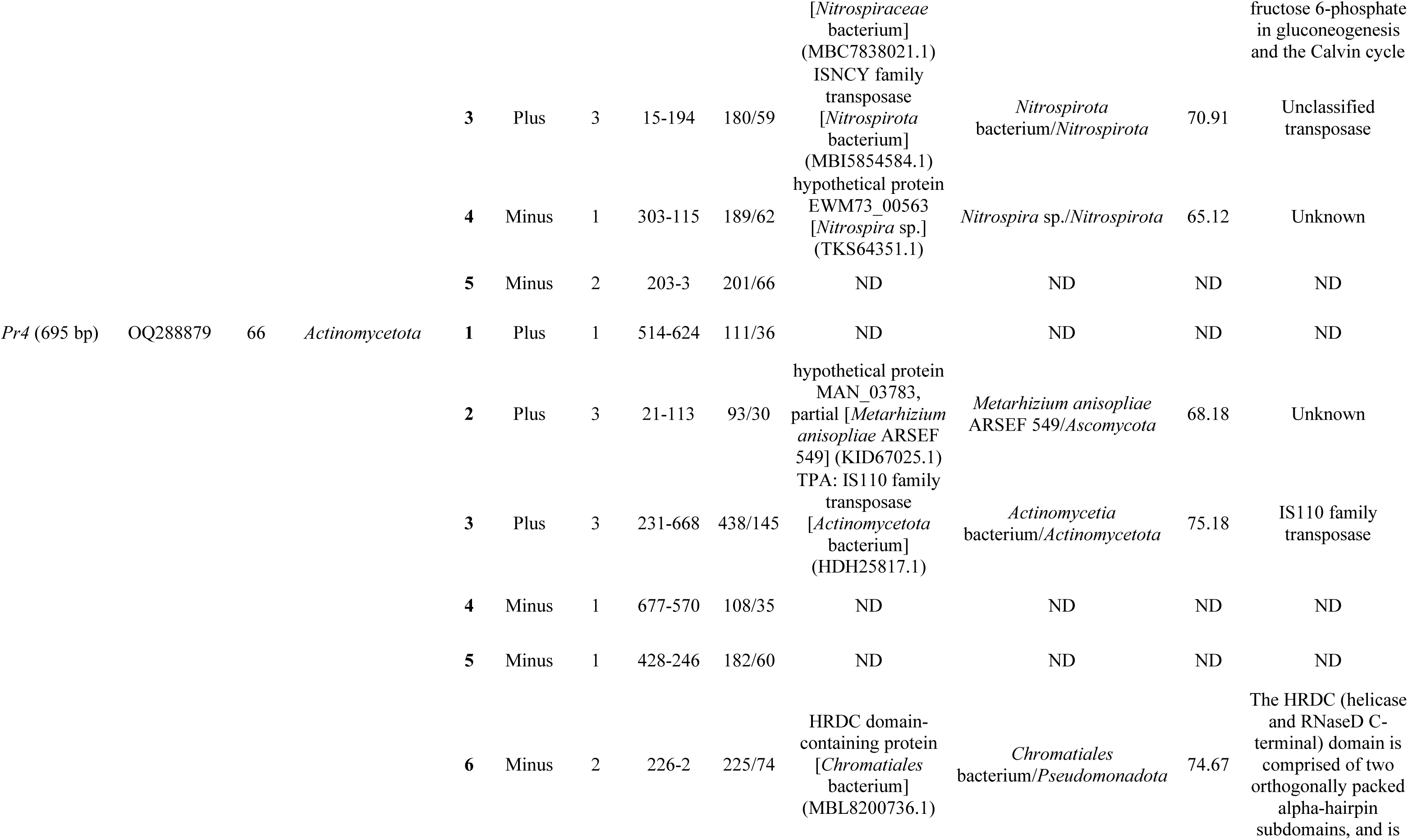
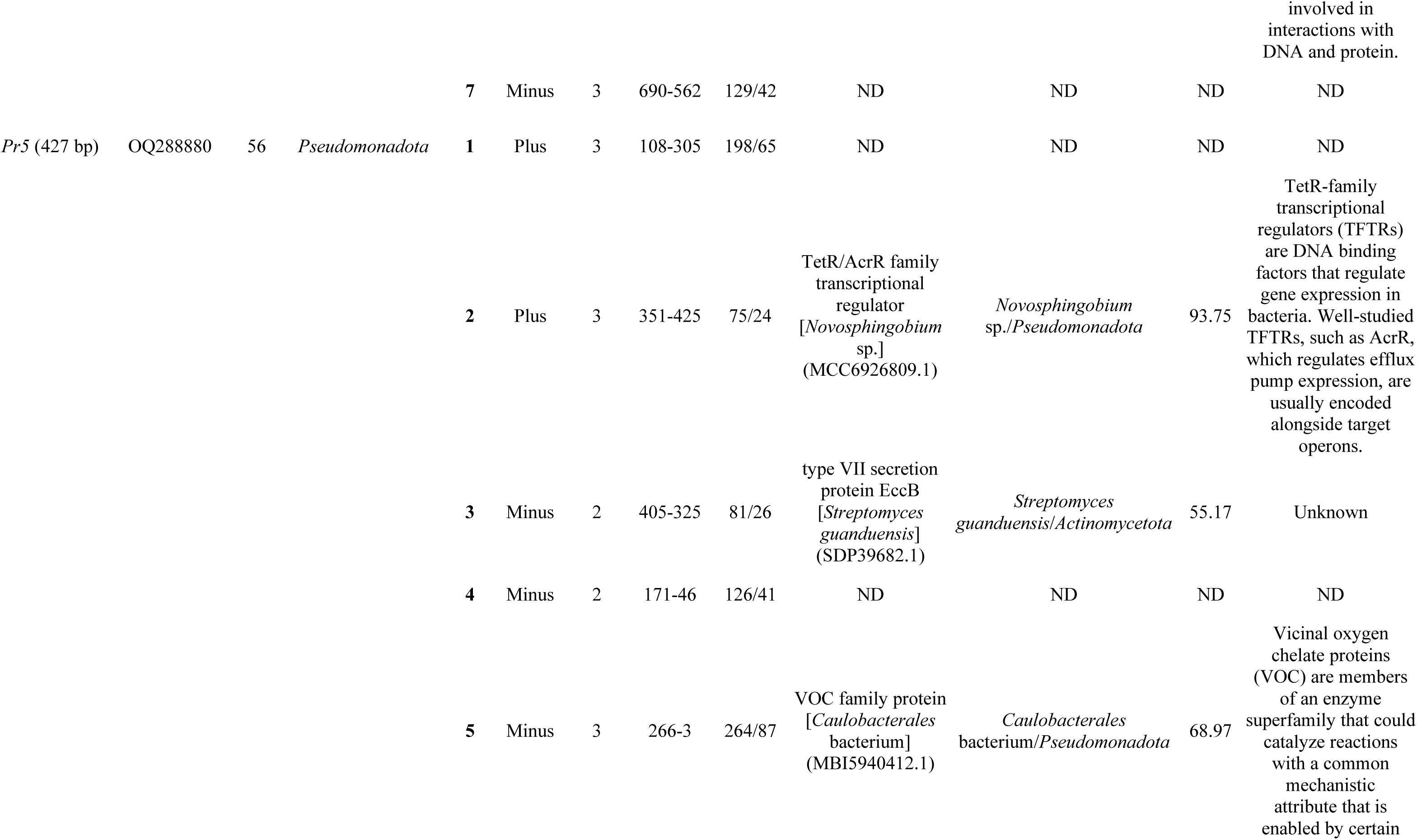
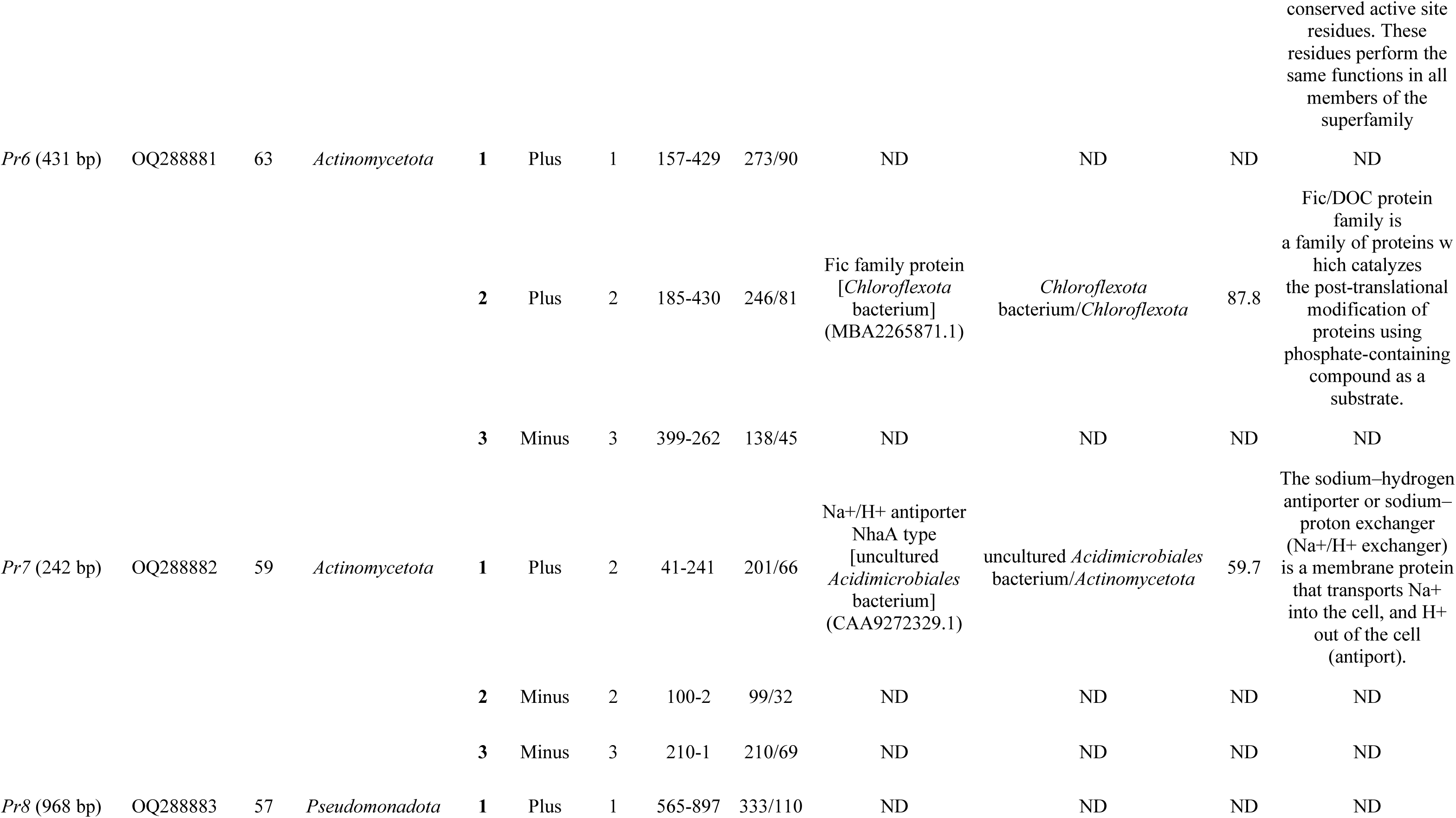
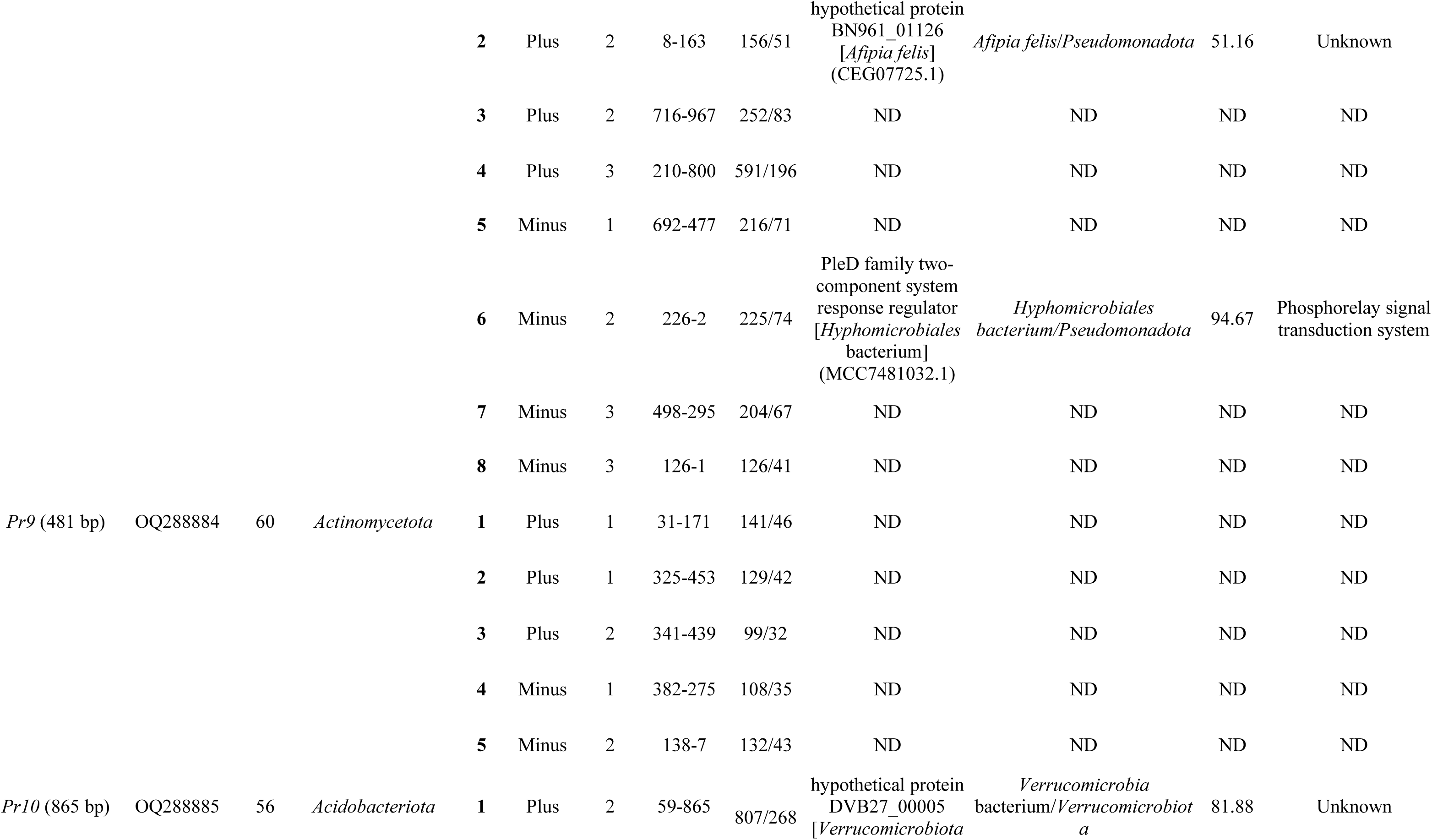
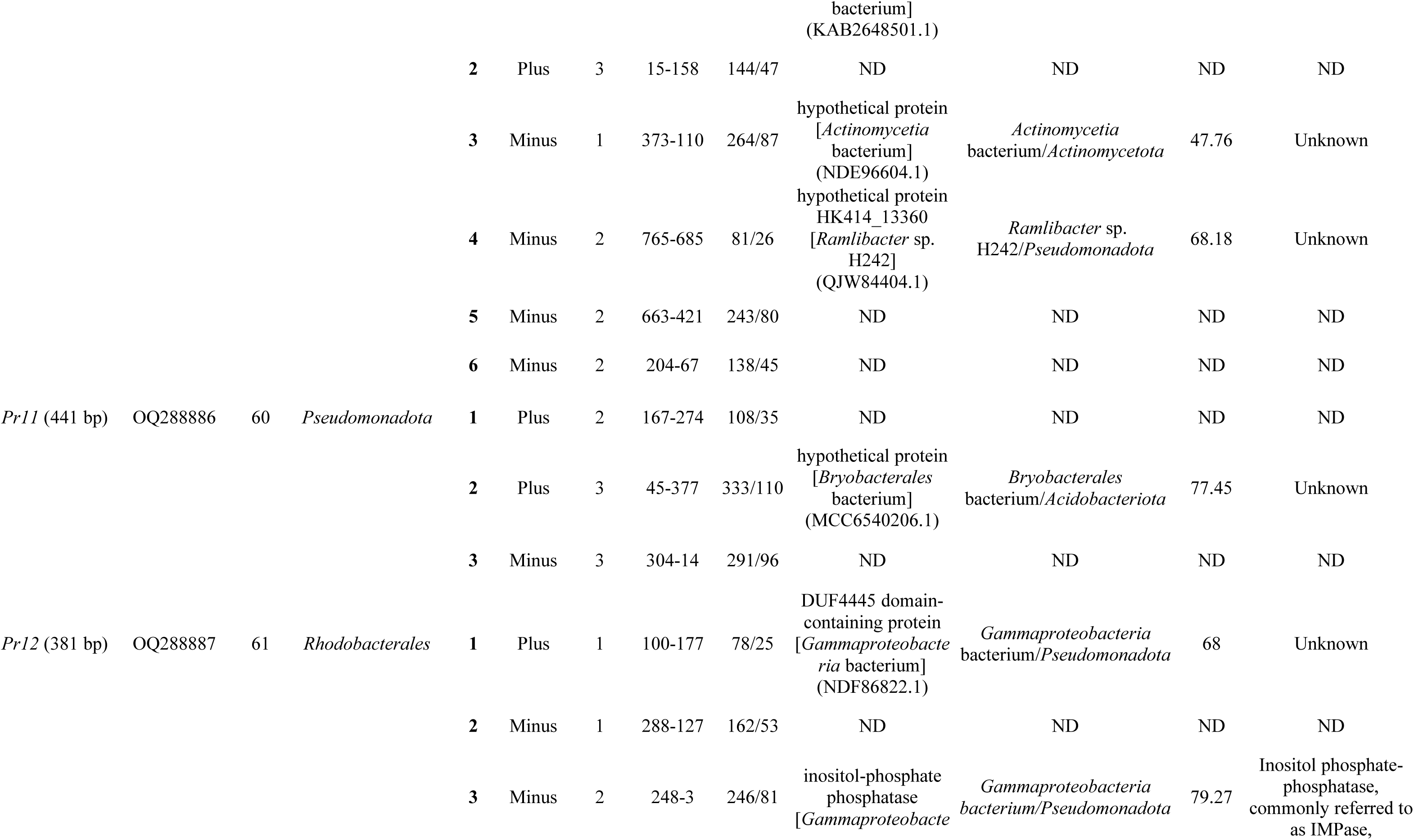
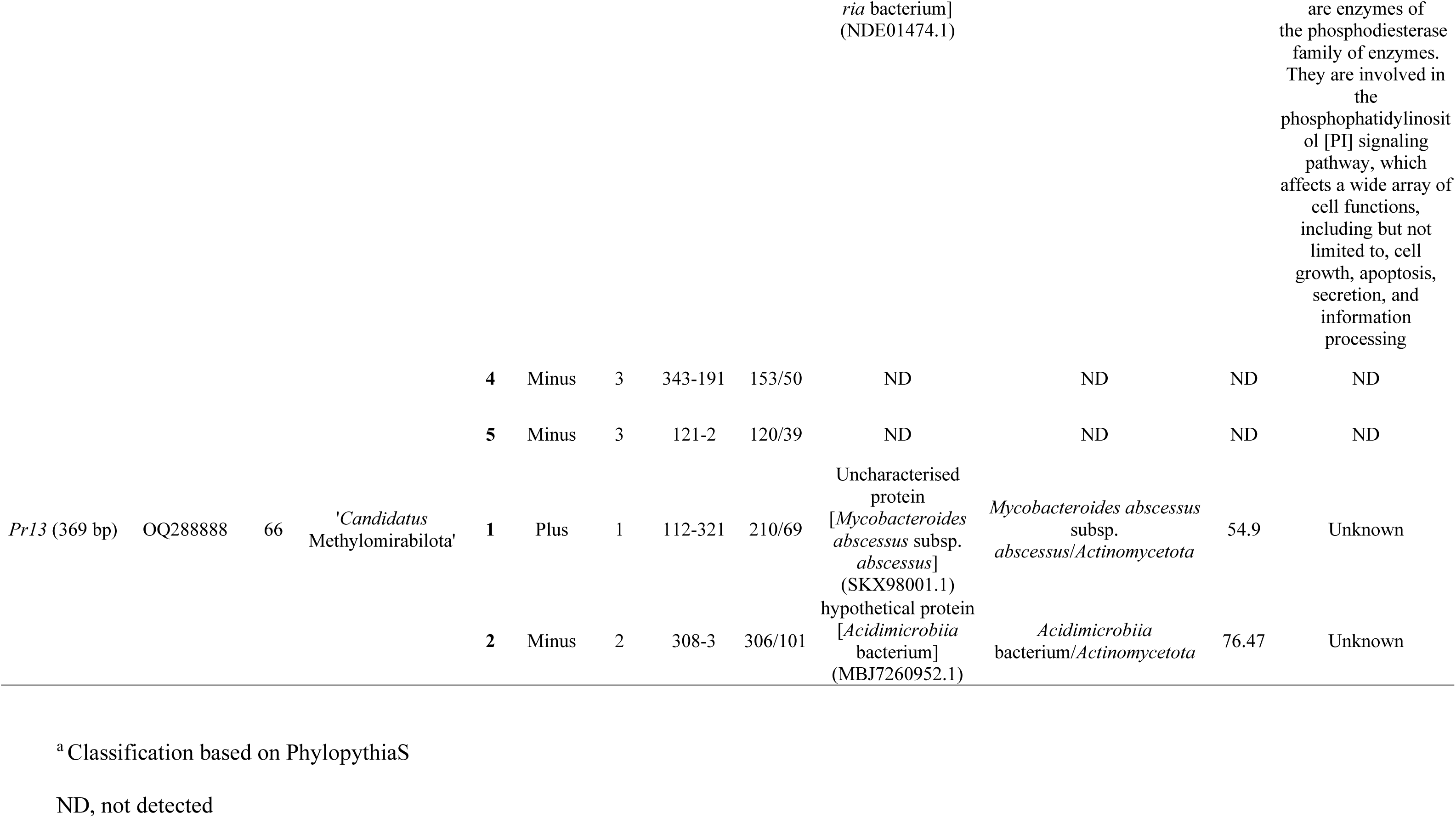
*In silico* characterization of the metagenomic DNA fragments with promoter activity.

Sixty ORFs were identify considering both strands of the insert sequences, ranging from eight ORFs (*Pr8*) to two ORFs (*Pr13*), with a mean of three ORFs per insert (Table 2). For each ORF, their putative function was determined by comparison of their amino acid sequence against the NCBI database. Interestingly, for more than half of the identified ORFs (53%) no significant similarity was found. For those ORFs with putative functions retrieved, four were of unknown function and the rest were associated with metabolic and regulatory functions.

### 3.3 Putative promoter sequences were predicted by in silico analysis

In order to determine putative promoter sequences present in the metagenomic inserts, three different software (BacPP, SAPPHIRE and BDGP) were used for *in silico* promoter prediction. The analysis retrieved 164 promoter sequences among the 13 inserts (Table S3), being 57, 86 and 21 putative promoters identified by BacPP, SAPPHIRE and BDGP, respectively. Figure 2 schematically represent the location of promoter sequences retrieved by the different software, regarding the identified ORFs and GFP coding sequence. As shown in Fig. 2, depending on the algorithm employed is the region detected as putative promoter. For some regions, there was a convergence between the predictions of the three algorithms (*e.g. Pr1A, Pr1B, Pr2A, Pr3A, Pr5B, Pr6A, Pr7A, Pr8C* and *Pr9A*), which suggest more confidence in those particular regions as putative promoter regions.

**Figure 2.**
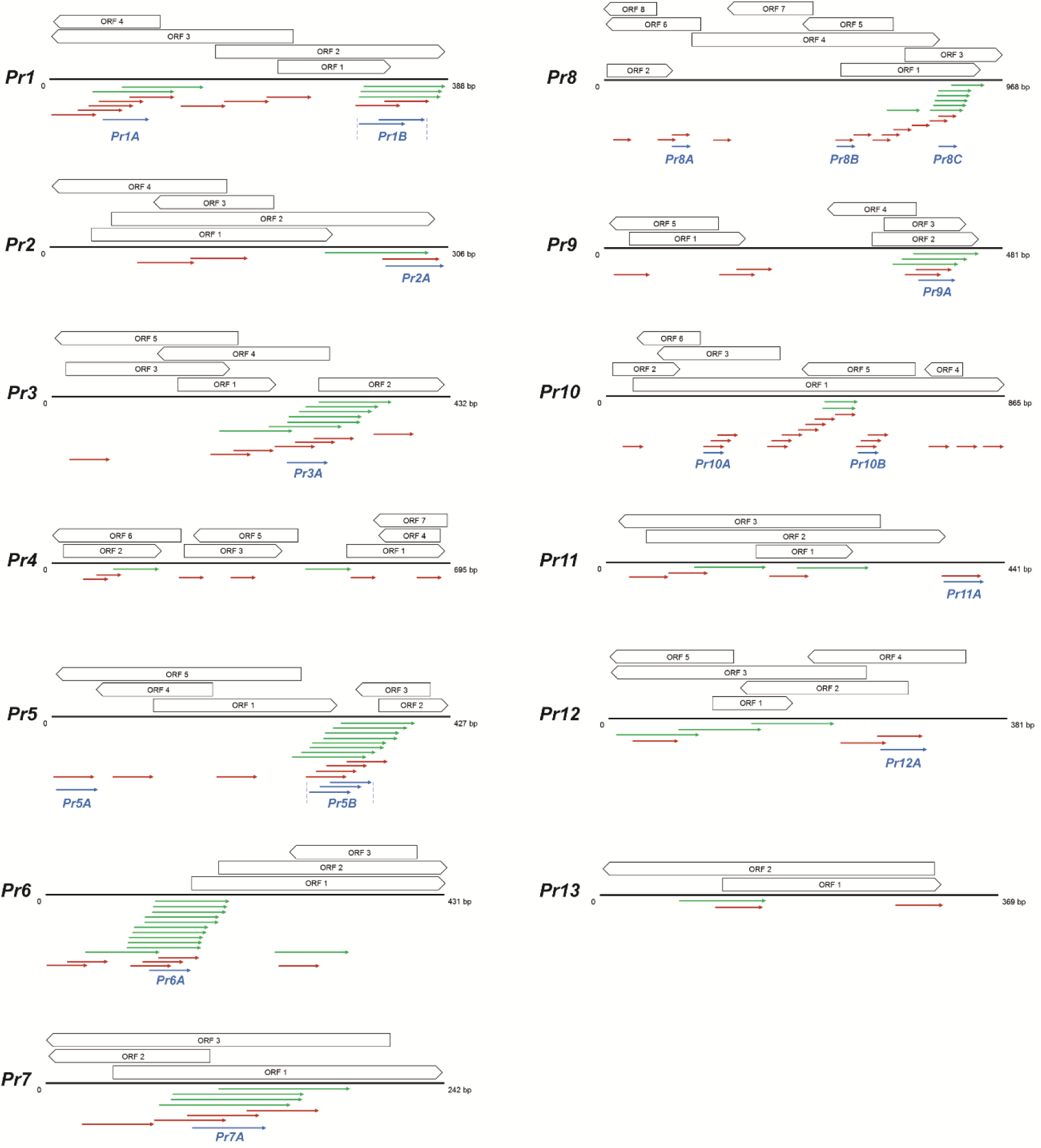
Schematic representation of the *in silico* analysis of the metagenomic DNA fragments with promoter activity. For each sequence (*Pr1-Pr13*), its length in bp and the location and orientation of the retrieved ORFs are shown. ORFs are numbered for each *Pr(x)* according to Table 2. For all the *Pr(x)*, *gfplva* gene is located on the right side end. Coloured arrows (BacPP, green; SAPPHIRE, red; BDGP, blue) indicate promoter sequences retrieved by the different algorithms. Details about predicted promoter sequences are in Table S3. All the features are at scale within a specific *Pr(x)*, but not between different *Pr(x)* sequences.

In order to determine if the retrieved promoter sequences harbor consensus motifs for the binding of the transcriptional machinery, first we identified these motifs in the promoter regions of *Pseudomonas* spp. As shown in Fig. S2A, a TTGN_3_-N_18_-TAN_3_T consensus motif was found in the promoter regions of *Pseudomonas* spp., TTGN_3_ corresponding to the −35 and TAN_3_T to −10, with a spacer of 18 nucleotides. These consensus motifs were found in promoter sequences retrieved by BDGP (Fig. S2B) but not in those retrieved by BacPP or SAPPHIRE (Fig. S2C and S2D, respectively).

Considering BDGP predicted the least amount of promoter sequences and that they were the only ones harboring the *Pseudomonas* spp. consensus motif, these promoters were selected for further analysis.

### 3.4 Genomic context, distance to gfp, and consensus motifs are all factors affecting promoter activity

As shown in Fig. 2, for some of the metagenomics inserts (*Pr1*, *Pr5*, *Pr8* and *Pr10*) BDGP (blue arrows) predicted more than one putative promoter sequence. In order to determine which of them is actually driving *gfp* expression in those inserts, BDGP predicted promoters *Pr3A* (50 bp), *Pr5A* (50 bp), *Pr5B* (71 bp), *Pr6A* (50 bp), *Pr8A* (50 bp), *Pr8B* (50 bp), *Pr8C* (50 bp), *Pr10A* (50 bp) and *Pr10B* (50 bp) were synthesized (Table S2) and their activity was evaluated with the same reporter system, as schematized in Fig. 3A. As shown in Fig. 3B, those promoters that in the metagenomic insert were closer to *gfp* (*Pr5B*, *Pr8C* and *Pr10B*) were the functional ones in all cases. Activity exerted by synthetic promoters *Pr5B* and *Pr10B*, was weaker than that of their respective complete metagenomic sequences (*Pr5* and *Pr10*). On the other hand, synthetic promoter *Pr8C* activity outperformed that exerted by complete sequence *Pr8*. These results suggest the presence of additional sequence elements in the original complete metagenomic fragment that play a role in promoter strength. Interestingly, the three active selected sequences (*Pr5B*, *Pr8C* and *Pr10B*) exerted the same promoter activity while their respective complete metagenomic sequences (*Pr5*, *Pr8* and *Pr10*) exerted activity of different strengths (Fig. 3B). This made us wonder if the observed behavior was related to having moved the promoters at the same distance to *gfp.* As proof of concept, we selected two additional promoter sequences to evaluate: *Pr3*, where promoter *Pr3A* is located at 295 bp to *gfp;* and *Pr6,* where promoter *Pr6A* is located at 440 bp to *gfp*. Both sequences were synthetized and were cloned at the same distance (159 bp) to *gfp* as *Pr5B, Pr8C* and *Pr10B* (Fig. 3A). As shown in Fig. 3B, when the promoter sequence *Pr3A* is moved closer to *gfp*, activity is the same than that of the selected active promoters *Pr5B, Pr8C* and *Pr10B,* while for selected sequence *Pr6A* activity was less. Therefore, promoter distance to *gpf* alone did not explain why the active selected promoters displayed the same promoter strength. Then, we analyze core promoter sequences −10 and −35 of all the selected promoters. In promoters *Pr3A*, *Pr5B*, *Pr8C* and *Pr10B* consensus promoter sequences of *Pseudomonas* spp. TTGN_3_ and TAN_3_T were conserved, while for *Pr6A* consensus sequences were less conserved (Table S4). We determined that selected promoters *Pr5A*, *Pr8A*, *Pr8B* and *Pr10A* (the inactive ones) also harbor less conserved consensus sequences than their respective active promoter sequences (*Pr5B*, *Pr8C* and *Pr10B*). Therefore, conservation of the consensus TTGN_3_ and TAN_3_T motifs are important for promoter strength.

**Figure 3.**
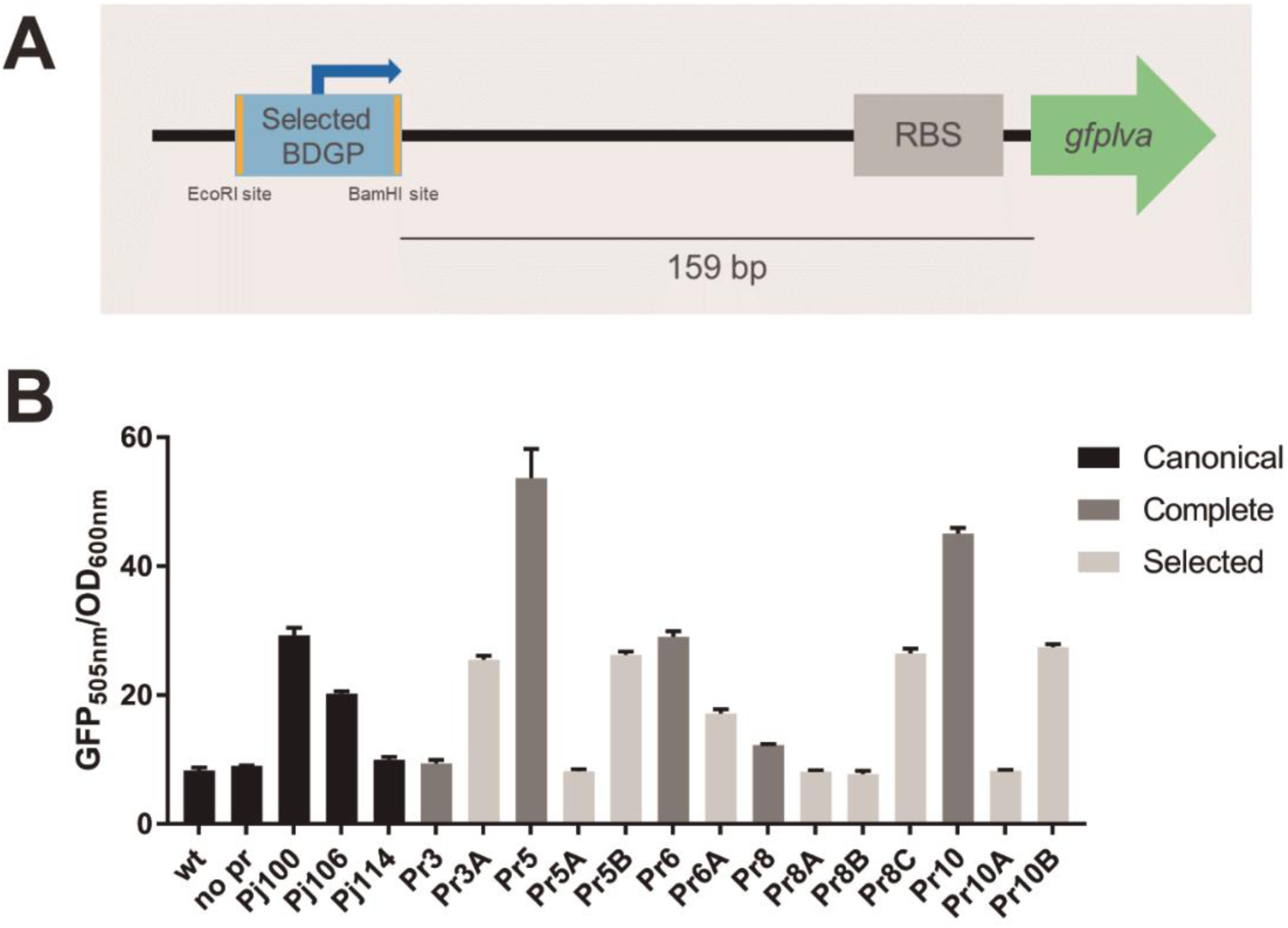
Promoter activity of selected promoters identified by BDGP in *P. putida* KT2440. **(A)** Schematic representation of the cloning site of the selected promoters (selected BDGP) in plasmid pSEVA231-*gfp*. **(B)** The functionality of the selected promoters *Pr3A*, *Pr3B*, *Pr5A*, *Pr5B*, *Pr6A*, *Pr8A*, *Pr8B*, *Pr8C*, *Pr10A*, and *Pr10B* was evaluated in *P. putida* KT2440. The parental strain (wt) and *P. putida* KT2440 harbouring promoter-trap plasmid without promoter (no pr) were used as negative controls. *P. putida* KT2440 harbouring plasmids with the Anderson promoters *Pj100*, *Pj106*, *Pj114* and the original metagenomic sequences with promoter activity (*Pr3*, *Pr5*, *Pr6*, *Pr8* and *Pr10)* were used for comparison. Expression was quantitative determined in cells grown in M9Km for 60 h, at 25 °C. Differences in GFP expression were evaluated by analysis of variance (ANOVA). The significances of the mean differences were estimated with the Tukey test (*p* < 0.05).

None of the selected promoter constructs generated a noteworthy metabolic burden to *P. putida* KT2440 as evidenced by the similar profiles of growth curves (Fig. S3).

### 3.5 Promoter sequences are functional in different Pseudomonadota

Despite the wide available repertoire of promoter sequences, most were identified and characterized in model microorganisms and are mostly host-specific. Broad host-range functional promoters are undoubtedly very interesting, as they can be used to drive expression in different chassis. With this in mind, we evaluated the performance of the promoter sequences in other *Pseudomonadota*. All 13 constructs (pSEVA231-*Pr(x)gfp*) were recovered from *P. putida* KT2440 and were introduced into Antarctic *Pseudomonas* sp. UYIF39, *E. coli* DH5α, *C. taiwanensis* R1^T^, *P. phymatum* STM 815^T^, and *E. meliloti* 1021.

As shown in Fig. 4, the metagenomics sequences exert different promoter activities depending on the host. In *Pseudomonas* sp. UYIF39 eleven constructs were functional. For this host, sequences *Pr2*, *Pr4*, *Pr5*, *Pr6*, and *Pr10* act as strong promoters similar to canonical promoter *Pj100*, while *Pr1*, *Pr7*, *Pr8*, *Pr12* and *Pr13* act as medium strength promoters if compared to canonical promoter *Pj106.* Sequence *Pr11* was functional in *Pseudomonas* sp. UYIF39 exerting a weak expression just above the controls, while sequences *Pr3* and *Pr9* were non-functional in this host (Fig. 4).

**Figure 4.**
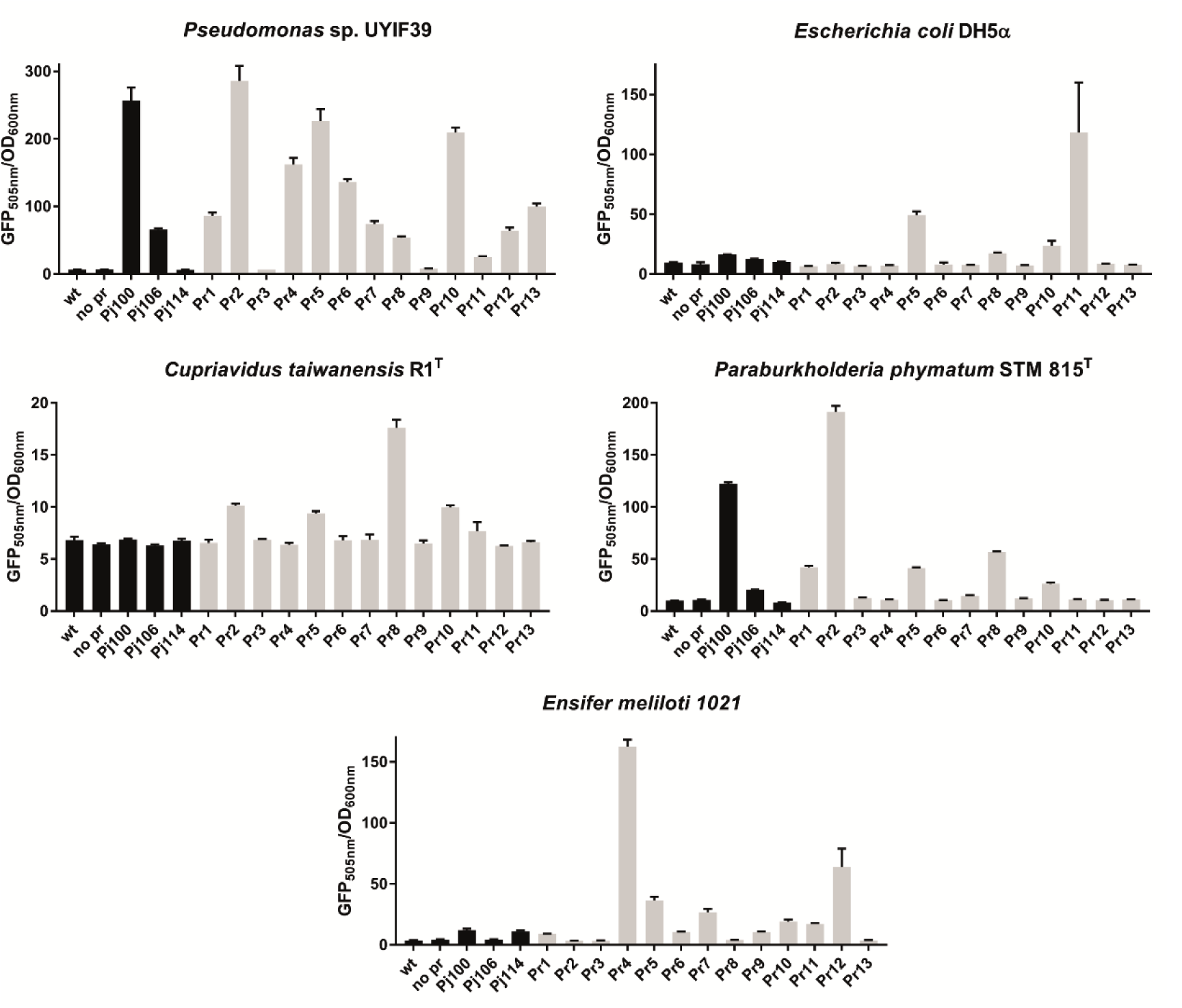
Activity of retrieved promoter sequences in different hosts. The functionality of the promoter sequences (*Pr1*-*Pr13*) was evaluated in different hosts belonging to the phylum *Pseudomonadota*: *Pseudomonas* sp. UYIF39, *E. coli* DH5α, *C. taiwanensis* R1^T^, *P. phymatum* STM 815^T^, and *E. meliloti* 1021. Parental strains (wt) and strains harbouring the promoter-trap plasmid without promoter (no pr) were used as negative controls. Strains harbouring Anderson promoters *Pj100*, *Pj106* and *Pj114* were used as controls to determine promoter strength. Quantitative GFP expression was determined in cells grown for 60 h in M9Km broth (M9SNm for *E. meliloti* 1021) at 25 °C. Fluorescence was normalized to OD_600nm_ for each strain and each construct. Results are the mean of three technical replicates in the same experiment. The experiment was repeated at least two times with similar results. Differences in GFP expression were evaluated by analysis of variance (ANOVA). The significances of the mean differences were estimated with the Tukey test (*p* < 0.05).

In *P. phymatum* STM 815^T^ five sequences were functional. Sequences *Pr1*, *Pr2*, *Pr5*, *Pr8* and *Pr10* were capable of driving *gfp* expression, acting *Pr2* as a stronger promoter than the canonical promoter *Pj100* (Fig. 4). In *C. taiwanensis* R1^T^, four sequences (*Pr2*, *Pr5*, *Pr8* and *Pr10*) exerted some *gfp* expression, while this host did not recognize any of the Anderson promoters. In *E. meliloti* 1021, sequences *Pr4*, *Pr5, Pr7, Pr10, Pr11* and *Pr12* exerted expression, being *Pr4* the strongest promoter. Anderson promoters were not recognized by *E. meliloti* and, surprisingly, this was also observed in *E. coli* DH5α, where only promoters *Pr5* and *Pr11* exerted expression (Fig. 4).

Regarding the fitness cost of the constructions and the exerted expression in the different host (Fig. S4-S8), slightly differences can be observed depending on the host and the construct they harbor. Differences are mainly related to longer lag phases and/or lower optical densities induced by the presence of some constructs.

The difference in the expression pattern of the *Pr(x)* sequences in the different host is most probably related to differences in transcriptional machinery and their recognition of promoter sequences. To gain insight in this regard, we evaluated the similarity among transcriptional factors σ^70^ between hosts. As shown in Table S5, the highest similarity with *P. putida* KT2440 σ^70^ was that of *Pseudomonas* sp. UYIF39, with an 87.34% sequence identity. This agrees with *Pseudomonas* sp. UYIF39 being the host where most of the sequences retrieved from *P. putida* KT2440 were functional. For the other strains, sequence identity was below 67%, which might explain the few *Pr(x)* sequences that were recognized by these hosts.

We can conclude that sequence *Pr5* is the most promising promoter sequence as it is functional in all the hosts assessed (Figure 5). Sequences *Pr2, Pr8,* and *Pr10* are functional in four of the six hosts (both *Pseudomonas*, *P. phymatum*, and *C. taiwanensis*), and *Pr4* is functional in both *Pseudomonas* and in *E. meliloti*.

**Figure 5.**
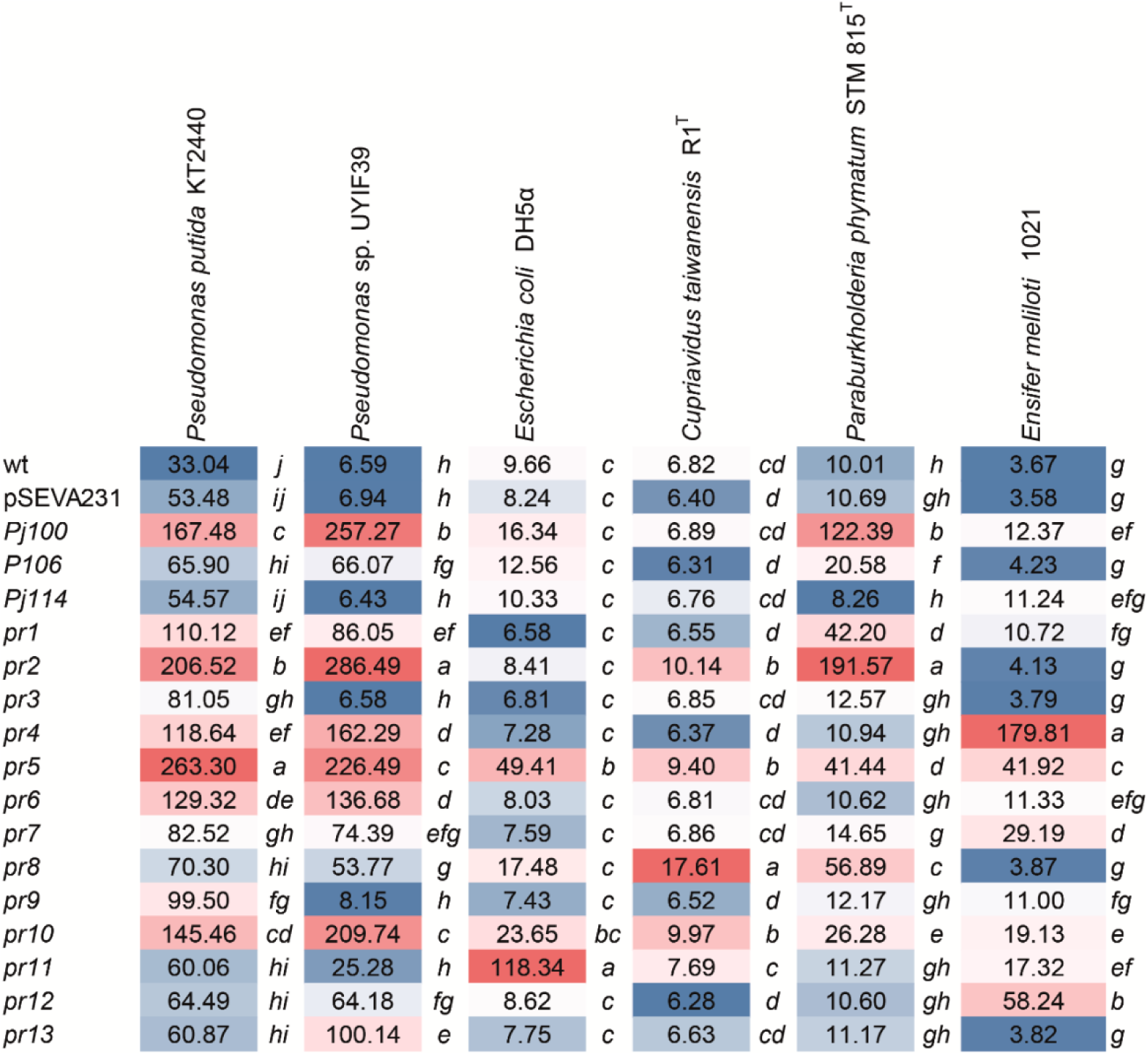
Heatmap of promoter activity in the different hosts. Promoter relative activity of each sequence (*Pr1-Pr13*) in each host is shown as a heatmap (red, highly expressed; blue, least expressed). Statistical analysis was performed independently for each host harbouring the different constructs (pSEVA231-*Pr*(*x*)*gfp*). Therefore, comparisons regarding promoter strength can only be made among constructs from the same genomic background. Different letters indicate significant differences between different promoters for the same host (Tukey, *p* < 0.05).

## 4. Discussion

Here we use a promoter-trap approach to find novel broad host-range promoter sequences functional in different *Pseudomonadota*. Using *gfp* as reporter, *Pseudomonas putida* KT2440 as chassis, and Antarctic soil metagenomic DNA as input, we generated a library of 1,804 clones and identified 13 unique functional promoter sequences.

The *in silico* analysis of the metagenomic fragments with promoter activity determined they were primarily associated with the phyla *Actinomycetota* and *Pseudomonadota* (Table 2). These results are in agreement with the high abundance of members of these phyla in samples from Fildes Peninsula, Antarctica, were the metagenomic DNA comes from ^27^. Additionally, these results suggest a bias in the ability of *P. putida* KT2440 transcriptional machinery to recognize sequences from those particular phyla, probably due to their GC content ^28^. Active metagenomic fragments were in the range of 56-66%, which is close to the genome GC content of *P. putida* KT2440 that has a mean of 61.6% ^29^. As a counterexample, no active metagenomics fragment associated with low GC phyla, such as *Bacillota*, were recovered. This could be due to functional aspects related to GC content, or to a low abundance of these phyla in our metagenomic sample. The abundance of *Bacillota* in Antarctic soils is variable, ranging from 3% to 62.2% depending on the sample ^30^. Therefore, the reason for the “no-*Bacillota*” bias in our approach should be elucidated by random sequencing of metagenomics inserts of non-active clones of the promoter-trap library. Despite interesting, was out of the scope of this work.

The different metagenomic fragments (*Pr1-Pr13*) exerted different promoter strength in *P. putida* KT2440 (Fig. 1) which is most likely related to differences in transcription, but we cannot rule out differences in translation or mRNA stabilization. Differences in transcription might be related to how conserved the consensus −35 and −10 sequences are ^31^, the length of the spacer between both motifs ^32^, and the presence of UP-elements (upstream promoter elements) ^33^, among other factors. Promoter sequences *Pr2A*, *Pr3A*, *Pr5B*, *Pr8C*, *Pr9A* and *Pr10B* predicted by BDGP, all harbor the *Pseudomonas* spp. promoter consensus sequences TTGN_3_ and TAN_3_T (Table S4). Despite this, no correlation between the presence of conserved consensus sequences and the promoter strength of the metagenomic inserts was found. Sequences *Pr2* and *Pr5* exerted strong promoter activity while sequences *Pr9* and *Pr10*, and *Pr3* and *Pr8*, exerted a medium-high and medium strength activity, respectively. This suggests that elements other than the consensus −35 and −10 are needed for promoter strength. This was also evident when comparing the expression profiles exerted by the whole inserts *vs* that exerted by their corresponding selected promoters (Fig. 3B). Selected promoters are synthetic 50bp sequences harboring only the promoter sequences retrieved by BDGP, therefore harboring the −35 and −10 boxes but lacking any additional elements present in their respective metagenomic insert that might be important for promoter strength. The fact that all selected sequences (with the exception of *Pr6A*) exerted the same promoter activity while their corresponding metagenomic inserts exerted different promoter strengths (Fig. 3B) suggests that the sequence context of the core promoter contain key elements that determines promoter strength. Further work is needed to elucidate this. Besides this, consensus sequences −35 and −10 also proved to be relevant for promoter strength as active selected promoters *Pr5B*, *Pr8C* and *Pr10B* harbor the consensus sequence TTGN_3_ and TAN_3_T, while the inactive ones (*Pr5A*, *Pr8A*, *Pr8B*, *Pr10A*) harbor somehow imperfect consensus (Table S4). The importance of these sequences was also evident in selected promoter *Pr6A,* which harbors an imperfect −10 box (Table S4) and exerted less promoter activity than that of the other active selected promoters (*Pr3A*, *Pr5B*, *Pr8C*, and *Pr10B*) (Fig. 3B) which harbor a perfect TAN_3_T (Table S4).

From the sixty ORFs identified *in silico*, only twenty-four of them displayed some sequence identity with proteins from databases (Table 2), which could be related with under-representation of rare samples, such as those from Antarctica, in databases. In that sense, for eighteen of the twenty-four predicted proteins, the highest similarity scores were with sequences retrieved from metagenome assembly genomes (MAGs) from rare samples such as groundwater, terrestrial subsurface samples, hydrothermal sediments, and soil crust from the Negev desert. Interestingly, many sequences were retrieved from cold samples such as Lille Firn glacier, Mackay Glaciers regions, and seasonally ice-covered lakes (Table 2).

Regarding the functionality of the promoters in different *Pseudomonadota*, the results were host-dependent (Fig. 4 and Fig. 5), highlighting the importance of using the host of interest when searching new functions by a function-driven metagenomic approach. Despite this, we retrieved sequence *Pr5* that proved to be a broad host-range promoter functional in representatives of classes α, β, and γ *Proteobacteria*. This makes *Pr5* a very promising broad host-range promoter for the SynBio toolkit. Promoters *Pr2, Pr8,* and *Pr10* are also very interesting to continue working, as they are functional in β and γ *Proteobacteria*, as well as *Pr4*, which is functional in α, and γ *Proteobacteria.* Sequence *Pr4* is particularly interesting as no canonical σ^70^ dependent promoter sequence was retrieved by BDGP, suggesting other promoter sequences are involved in expression.

The importance of having broad host-range promoters, or promoters specifically designed for the host of interest, is also evidenced by the behavior of Anderson promoters in the different host (Fig. 5). These promoter sequences, as most of the promoter sequences available, are designed for *E. coli* ^20^. The promoters addressed here (*Pj100*, *Pj106* and *Pj114*) were nonfunctional neither in *C. taiwanensis* nor in *E. meliloti* (Fig. 5). For *E. meliloti*, there is only one report from an iGEM team who used *Pj100* fused to citrine, showing some expression but only 1.5x above the control strain ^34^. Regarding *C. taiwanensis,* to the best of our knowledge, this is the first report of Anderson promoters being assessed in this host. However, in a recent work promoters *Pj100* and *Pj106* were evaluated in *Cupriavidus necator* and proved to be functional ^7^. This suggests a species-dependent behavior of these promoters. Remarkably, Anderson promoters were not functional in *E. coli* DH5α (Fig. 5), which was not an expected outcome. This phenomena might be related to a strain-dependent behavior, as some iGEM teams reported differences in the promoter strength of Anderson promoters when assessed in *E. coli* DH5α ^35,36^, regarding the original results. Particularly, the performance of *Pj100* in *E. coli* strains BL21 (DE3), TOP10, and DH5α, was evaluated being promoter activity in DH5α significantly lower compared with the other two strains ^36,37^. Jervis *et al.* also used Anderson promoters in *E. coli* DH5α reporting a 2-fold reduction in promoter strength of J23113 regarding J23110, while for the original description a 100-fold reduction is reported ^38^. Recently, Pearson *et al.* ^39^ evaluated all Anderson promoters in *E. coli* XL1-blue, finding also some discrepancies to the original description. Considering the same example from above, expression from J23113 was higher (1.2-fold increase) to that from J23100 ^40^. Therefore, evaluating the performance of Anderson promoters in different strains of *E. coli* with the same reporter construction is very interesting and will be addressed by our group in a future work.

In summary, we identify and characterize a set of broad host-range promoters that will be of great use in SynBio. We also increased the repertoire of constitutive promoters functional in *P. putida* KT2440, the most promising novel chassis in the field. We can conclude that function-driven metagenomics is an excellent and rapid approach for searching new broad host-range promoters and that the Antarctic microbial community is a valuable resource for prospecting new biological parts for SynBio. In future work, we will continue characterizing the potential of the broad-host range promoters by evaluating new hosts and by rational/random sequence variations in order to increase or decrease the promoter strength.

## Author Contributions

VA conceived the study and supervised the experimental work. DMR performed the experiments. VA and DMR wrote the manuscript.

## Supplementary Material

Additional results regarding strain fitness, consensus promoter motifs, and promoter sequences retrieved by different algorithms are in supplementary tables and figures, available with the online version of this article.

## Acknowledgments

This work was partially funded by the Agencia Nacional de Investigación e Innovación (ANII, FCE_3_2018_1_148885), and PEDECIBA (Programa de Desarrollo de las Ciencias Básicas).

## References

(1) Promoters/Catalog. https://parts.igem.org/Promoters/Catalog.

(2) Promoters/Catalog/Anderson. http://parts.igem.org/Promoters/Catalog/Anderson.

(3) Calero, P.; Nikel, P. I. Chasing Bacterial Chassis for Metabolic Engineering: A Perspective Review from Classical to Non-traditional Microorganisms. Microb. Biotechnol. 2019, 12 (1), 98–124. 10.1111/1751-7915.13292.

(4) Schuster, L. A.; Reisch, C. R. A Plasmid Toolbox for Controlled Gene Expression across the Proteobacteria. Nucleic Acids Res. 2021, 49 (12), 7189–7202. 10.1093/nar/gkab496.

(5) Schuster, L. A.; Reisch, C. R. A Plasmid Toolbox for Controlled Gene Expression across the Proteobacteria. Nucleic Acids Res. 2021, 49 (12), 7189–7202. 10.1093/nar/gkab496.

(6) Yang, S.; Liu, Q.; Zhang, Y.; Du, G.; Chen, J.; Kang, Z. Construction and Characterization of Broad-Spectrum Promoters for Synthetic Biology. ACS Synth. Biol. 2018, 7 (1), 287–291. 10.1021/acssynbio.7b00258.

(7) Keating, K. W.; Young, E. M. Systematic Part Transfer by Extending a Modular Toolkit to Diverse Bacteria. ACS Synth. Biol. 2023, 12 (7), 2061–2072. 10.1021/acssynbio.3c00104.

(8) Taupp, M.; Mewis, K.; Hallam, S. J. The Art and Design of Functional Metagenomic Screens. Curr. Opin. Biotechnol. 2011, 22 (3), 465–472. 10.1016/j.copbio.2011.02.010.

(9) Westmann, C. A.; Alves, L. de F.; Silva-Rocha, R.; Guazzaroni, M.-E. Mining Novel Constitutive Promoter Elements in Soil Metagenomic Libraries in Escherichia Coli. Front. Microbiol. 2018, 9. 10.3389/fmicb.2018.01344.

(10) Lee, S. H.; Kim, J. M.; Lee, H. J.; Jeon, C. O. Screening of Promoters from Rhizosphere Metagenomic DNA Using a Promoter-trap Vector and Flow Cytometric Cell Sorting. J. Basic Microbiol. 2011, 51 (1), 52–60. 10.1002/jobm.201000291.

(11) Han, S.-S.; Lee, J.-Y.; Kim, W.-H.; Shin, H.-J.; Kim, G.-J. Screening of Promoters from Metagenomic DNA and Their Use for the Construction of Expression Vectors. J. Microbiol. Biotechnol. 2008, 18 (10), 1634–1640.

(12) Amarelle, V.; Sanches-Medeiros, A.; Silva-Rocha, R.; Guazzaroni, M.-E. Expanding the Toolbox of Broad Host-Range Transcriptional Terminators for Proteobacteria through Metagenomics. ACS Synth. Biol. 2019, 8 (4), 647–654. 10.1021/acssynbio.8b00507.

(13) Martínez-García, E.; de Lorenzo, V. Pseudomonas Putida as a Synthetic Biology Chassis and a Metabolic Engineering Platform. Curr. Opin. Biotechnol. 2024, 85, 103025. 10.1016/j.copbio.2023.103025.

(14) Martin-Pascual, M.; Batianis, C.; Bruinsma, L.; Asin-Garcia, E.; Garcia-Morales, L.; Weusthuis, R. A.; van Kranenburg, R.; Martins dos Santos, V. A. P. A Navigation Guide of Synthetic Biology Tools for Pseudomonas Putida. Biotechnol. Adv. 2021, 49, 107732. 10.1016/j.biotechadv.2021.107732.

(15) Zobel, S.; Benedetti, I.; Eisenbach, L.; de Lorenzo, V.; Wierckx, N.; Blank, L. M. Tn7-Based Device for Calibrated Heterologous Gene Expression in Pseudomonas Putida. ACS Synth. Biol. 2015, 4 (12), 1341–1351. 10.1021/acssynbio.5b00058.

(16) Elmore, J. R.; Furches, A.; Wolff, G. N.; Gorday, K.; Guss, A. M. Development of a High Efficiency Integration System and Promoter Library for Rapid Modification of Pseudomonas Putida KT2440. Metab. Eng. Commun. 2017, 5, 1–8. 10.1016/j.meteno.2017.04.001.

(17) Amarelle, V.; Roldán, D. M.; Fabiano, E.; Guazzaroni, M.-E. Synthetic Biology Toolbox for Antarctic Pseudomonas Sp. Strains: Toward a Psychrophilic Nonmodel Chassis for Function-Driven Metagenomics. ACS Synth. Biol. 2023. 10.1021/acssynbio.2c00543.

(18) Sambrook, J.; Fritsch, E. F.; Maniatis, T. Molecular Cloning-A Laboratory Manual; Cold Spring Harbor Laboratory: New York, USA, 1989.

(19) Figurski, D. H.; Helinski, D. R. Replication of an Origin-Containing Derivative of Plasmid RK2 Dependent on a Plasmid Function Provided in Trans. Proc. Natl. Acad. Sci. 1979, 76 (4), 1648–1652. 10.1073/pnas.76.4.1648.

(20) Promoters/Catalog/Anderson.

(21) Patil, K. R.; Roune, L.; McHardy, A. C. The PhyloPythiaS Web Server for Taxonomic Assignment of Metagenome Sequences. PLoS One 2012, 7 (6), e38581. 10.1371/journal.pone.0038581.

(22) de Avila e Silva, S.; Echeverrigaray, S.; Gerhardt, G. J. L. BacPP: Bacterial Promoter Prediction—A Tool for Accurate Sigma-Factor Specific Assignment in Enterobacteria. J. Theor. Biol. 2011, 287, 92–99. 10.1016/j.jtbi.2011.07.017.

(23) Coppens, L.; Lavigne, R. SAPPHIRE: A Neural Network Based Classifier for Σ70 Promoter Prediction in Pseudomonas. BMC Bioinformatics 2020, 21 (1), 415. 10.1186/s12859-020-03730-z.

(24) Reese, M. G. Application of a Time-Delay Neural Network to Promoter Annotation in the Drosophila Melanogaster Genome. Comput. Chem. 2001, 26 (1), 51–56. 10.1016/S0097-8485(01)00099-7.

(25) Aziz, R. K.; Bartels, D.; Best, A.; DeJongh, M.; Disz, T.; Edwards, R. A.; Formsma, K.; Gerdes, S.; Glass, E. M.; Kubal, M.; Meyer, F.; Olsen, G. J.; Olson, R.; Osterman, A. L.; Overbeek, R. A.; McNeil, L. K.; Paarmann, D.; Paczian, T.; Parrello, B.; Pusch, G. D.; Reich, C.; Stevens, R.; Vassieva, O.; Vonstein, V.; Wilke, A.; Zagnitko, O. The RAST Server: Rapid Annotations Using Subsystems Technology. BMC Genomics 2008, 9 (75), 1–15.

(26) Raes, J.; Korbel, J. O.; Lercher, M. J.; von Mering, C.; Bork, P. Prediction of Effective Genome Size in Metagenomic Samples. Genome Biol. 2007, 8 (R10), 1–11. 10.1186/gb-2007-8-1-r10.

(27) Cui, S.; Du, J.; Zhu, L.; Xin, D.; Xin, Y.; Zhang, J. Analysis of Microbial Diversity in South Shetland Islands and Antarctic Peninsula Soils Based on Illumina High-Throughput Sequencing and Cultivation-Dependent Techniques. Microorganisms 2023, 11 (10), 2517. 10.3390/microorganisms11102517.

(28) Gabor, E. M.; Alkema, W. B. L.; Janssen, D. B. Quantifying the Accessibility of the Metagenome by Random Expression Cloning Techniques. Environ. Microbiol. 2004, 6 (9), 879–886. 10.1111/j.1462-2920.2004.00640.x.

(29) Weinel, C.; Nelson, K. E.; Tümmler, B. Global Features of the Pseudomonas Putida KT2440 Genome Sequence. Environ. Microbiol. 2002, 4 (12), 809–818. 10.1046/j.1462-2920.2002.00331.x.

(30) Ramos, L. R.; Vollú, R. E.; Jurelevicius, D.; Rosado, A. S.; Seldin, L. Firmicutes in Different Soils of Admiralty Bay, King George Island, Antarctica. Polar Biol. 2019, 42 (12), 2219–2226. 10.1007/s00300-019-02596-z.

(31) Shultzaberger, R. K.; Chen, Z.; Lewis, K. A.; Schneider, T. D. Anatomy of Escherichia Coli σ 70 Promoters. Nucleic Acids Res. 2007, 35 (3), 771–788. 10.1093/nar/gkl956.

(32) Klein, C. A.; Teufel, M.; Weile, C. J.; Sobetzko, P. The Bacterial Promoter Spacer Modulates Promoter Strength and Timing by Length, TG-Motifs and DNA Supercoiling Sensitivity. Sci. Rep. 2021, 11 (1), 24399. 10.1038/s41598-021-03817-4.

(33) Ross, W.; Gosink, K. K.; Salomon, J.; Igarashi, K.; Zou, C.; Ishihama, A.; Severinov, K.; Gourse, R. L. A Third Recognition Element in Bacterial Promoters: DNA Binding by the α Subunit of RNA Polymerase. Science (80-. ). 1993, 262 (5138), 1407–1413. 10.1126/science.8248780.

(34) http://parts.igem.org/Part:BBa_K1856004.

(35) https://pubmed.ncbi.nlm.nih.gov/37294017/

(36) http://parts.igem.org/Part:BBa_J23100

(37) http://parts.igem.org/Part:BBa_J23100:Experience.

(38) Jervis, A. J.; Carbonell, P.; Taylor, S.; Sung, R.; Dunstan, M. S.; Robinson, C. J.; Breitling, R.; Takano, E.; Scrutton, N. S. SelProm: A Queryable and Predictive Expression Vector Selection Tool for Escherichia Coli. ACS Synth. Biol. 2019, 8 (7), 1478–1483. 10.1021/acssynbio.8b00399.

(39) Pearson, A. N.; Thompson, M. G.; Kirkpatrick, L. D.; Ho, C.; Vuu, K. M.; Waldburger, L. M.; Keasling, J. D.; Shih, P. M. The PGinger Family of Expression Plasmids. Microbiol. Spectr. 2023, 11 (3). 10.1128/spectrum.00373-23.

(40) Pearson, A. N.; Thompson, M. G.; Kirkpatrick, L. D.; Ho, C.; Vuu, K. M.; Waldburger, L. M.; Keasling, J. D.; Shih, P. M. The PGinger Family of Expression Plasmids. Microbiol. Spectr. 2023, 11 (3). 10.1128/spectrum.00373-23.

