## Supplementary material for "Identification of novel broad host-range promoter sequences functional in diverse *Pseudomonadota* by a promoter-trap approach"

[a] Departamento de Bioquímica y Genómica Microbianas. Instituto de Investigaciones Biológicas Clemente Estable. Av. Italia 3318, Montevideo 11600, Uruguay.

### Supporting Information

| | |
|---|---|
| **Figure S1.** Effect of plasmids in *P. putida* KT2440 fitness | S2 |
| **Figure S2**. Logo pattern obtained for promoter sequences | S3 |
| **Figure S3**. Effect of plasmids with selected promoter identified by BDGP in *P. putida* KT2440 fitness | S4 |
| **Figure S4**. Effect of plasmids in Antarctic *Pseudomonas* sp. UYIF39 fitness | S5 |
| **Figure S5**. Effect of plasmids in *E. coli* DH5α fitness | S6 |
| **Figure S6**. Effect of plasmids in *C. taiwanensis* R1T fitness | S7 |
| **Figure S7**. Effect of plasmids in *P. phymatum* STM 815T fitness | S8 |
| **Figure S8**. Effect of plasmids in *E. meliloti* 1021 fitness | S9 |
| **Table S1.** Strains and plasmids | S10 |
| **Table S2**. Primers used for the construction of the selected promoters | S13 |
| **Table S3**. Promoter sequences identified *in silico* within the metagenomic DNA fragments cloned in the promoter-trap vector | S14 |
| **Table S4**. DNA sequences of the active metagenomic fragments showing promoters predicted by BDGP and their features | S31 |
| **Table S5**. Similarity in the amino acid sequences of the transcription factor $\sigma^{70}$ or RpoD in the genomes of the strains used | S34 |
| References | S35 |

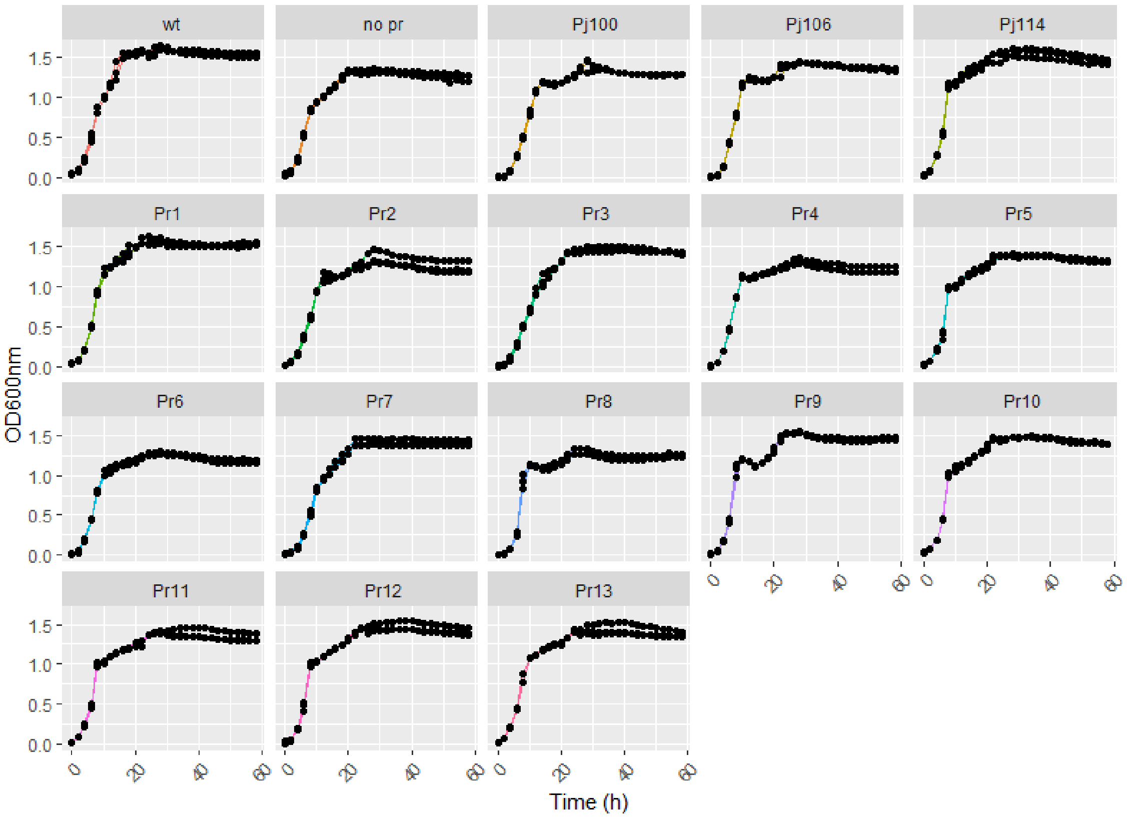

**Figure S1. Effect of plasmids in *P. putida* KT2440 fitness.** The effect on growth exerted by plasmids harboring the different promoter sequences and the different canonical promoters was evaluated. Strains were gown in LBKm at 25 °C for 16 h, diluted 20-fold in M9Km, and growth was evaluated at 25 °C every 2 h for a lapse of 60 h. Results are the mean of three technical replicates in the same experiment.

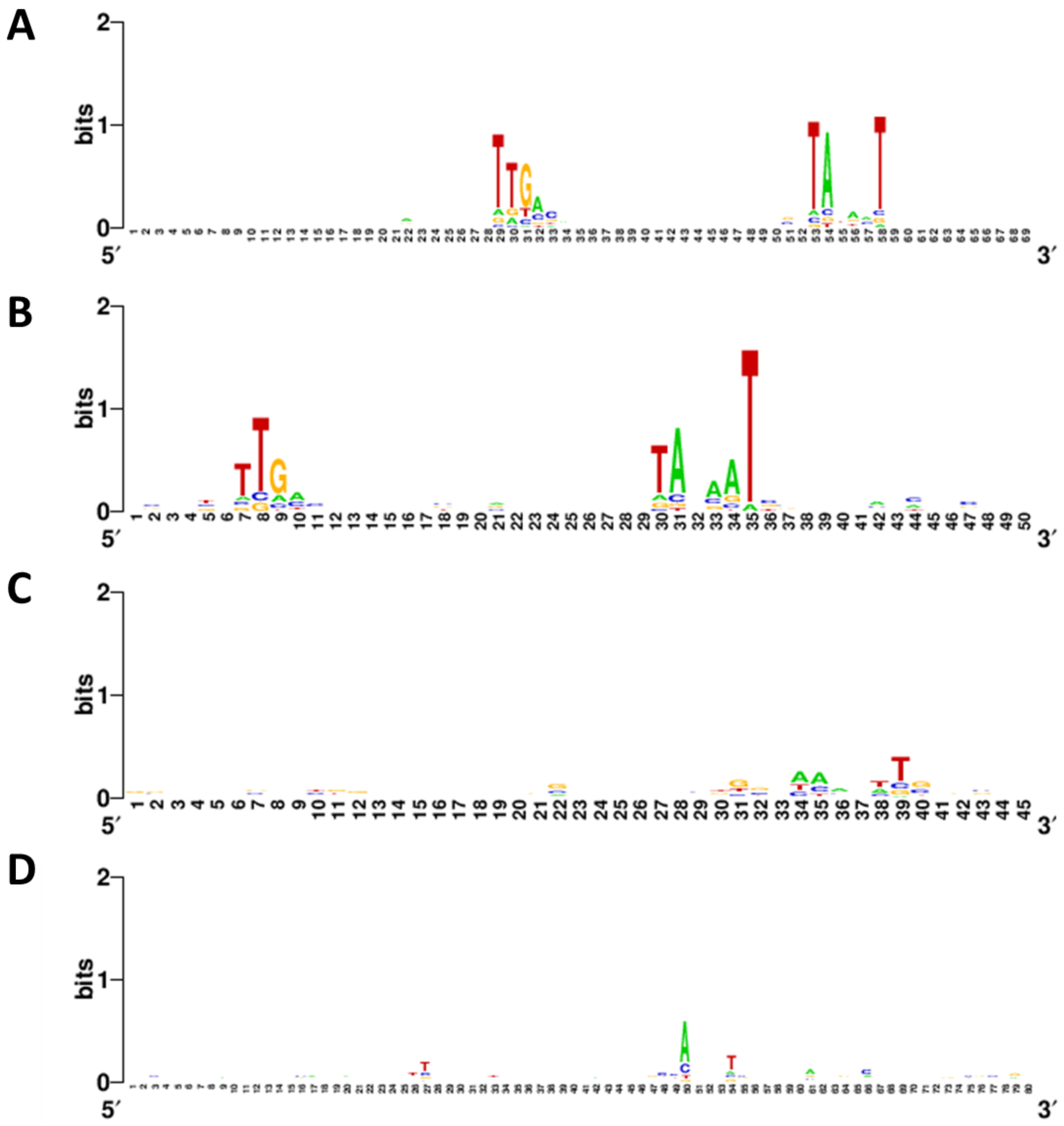

**Figure S2. Logo pattern obtained for promoter sequences.** Logo patterns were generated with WebLogo (Crooks et al. 2004). **(A)** Consensus sequence of 168 RpoD-dependent promoters of *Pseudomonas* spp. obtained from the SAPPHIRE database. Logo pattern of promoter sequences present in metagenomic DNA inserts, identified by the BDGP **(B)**, SAPPHIRE **(C)**, and BacPP **(D).**

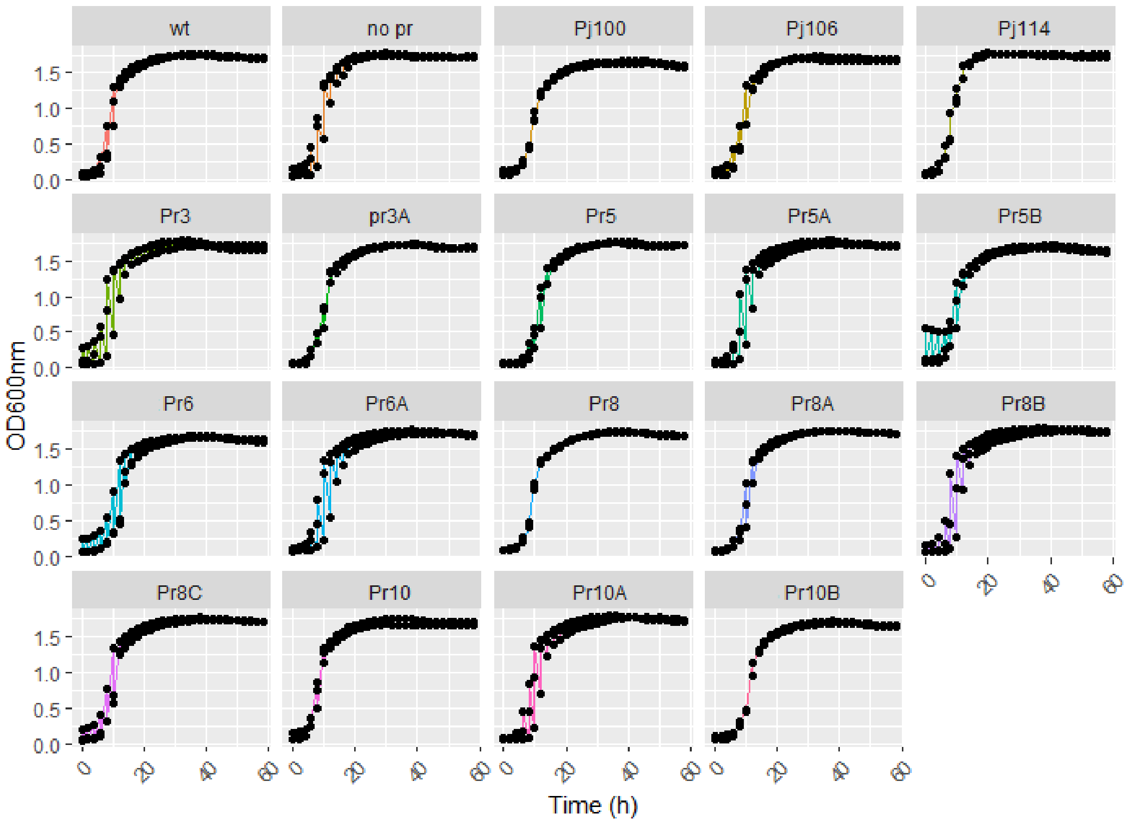

**Figure S3. Effect of plasmids with selected promoter identified by BDGP in *P. putida* KT2440 fitness.** The effect on growth exerted by plasmids harboring the different promoter sequences and the different canonical promoters was evaluated. Strains were gown in LBKm at 25 °C for 16 h, diluted 20-fold in M9Km, and growth was evaluated at 25 °C every 2 h for a lapse of 60 h. Results are the mean of three technical replicates in the same experiment.

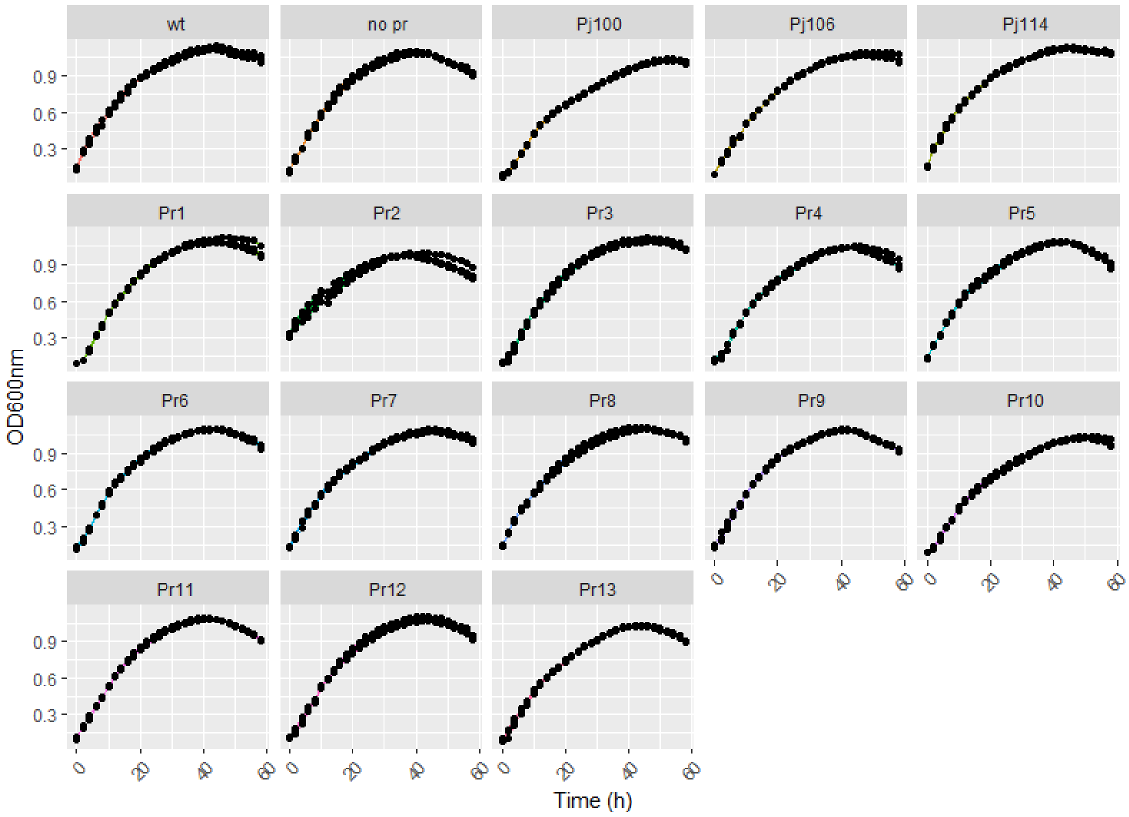

**Figure S4. Effect of plasmids in Antarctic *Pseudomonas* sp. UYIF39 fitness.** The effect on growth exerted by plasmids harboring the different promoter sequences and the different canonical promoters was evaluated. Strains were gown in LBKm at 25 °C for 16 h, diluted 20-fold in M9Km, and growth was evaluated at 25 °C every 2 h for a lapse of 60 h. Results are the mean of three technical replicates in the same experiment.

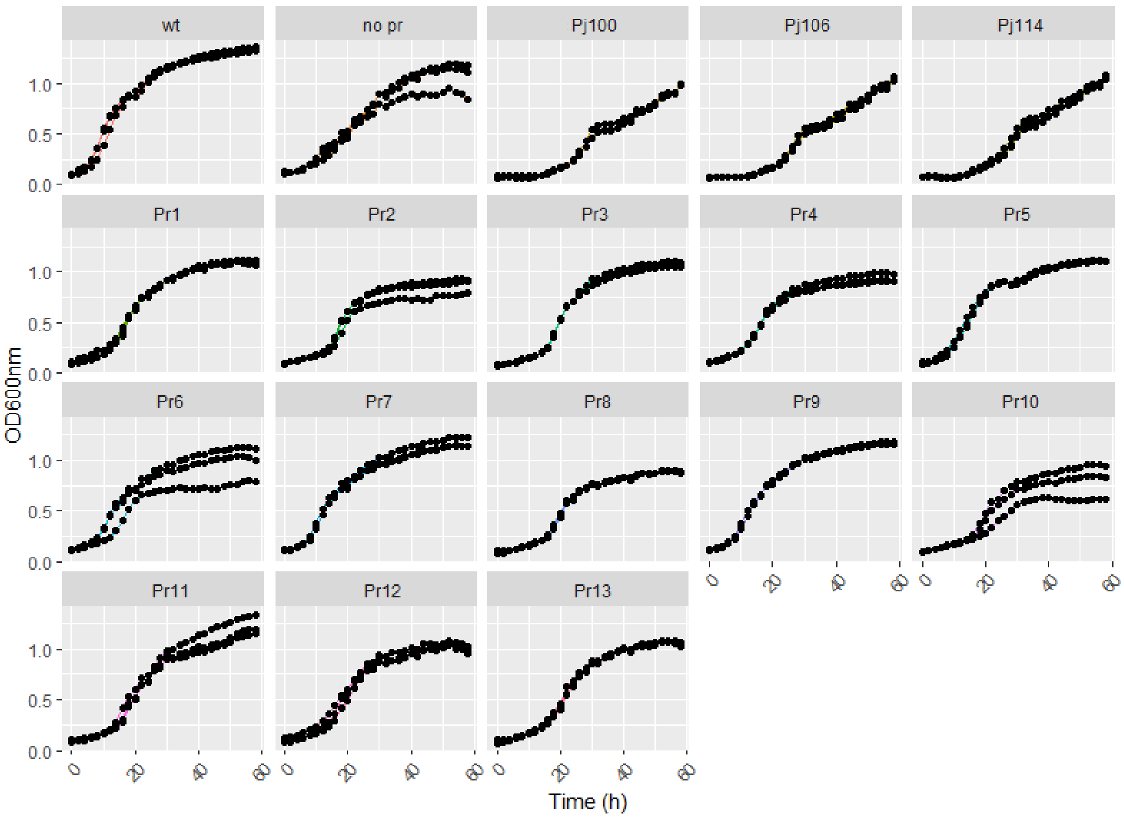

**Figure S5. Effect of plasmids in *E. coli* DH5α fitness.** The effect on growth exerted by plasmids harboring the different promoter sequences and the different canonical promoters was evaluated. Strains were gown in LBKm at 25 °C for 16 h, diluted 20-fold in M9Km, and growth was evaluated at 25 °C every 2 h for a lapse of 60 h. Results are the mean of three technical replicates in the same experiment.

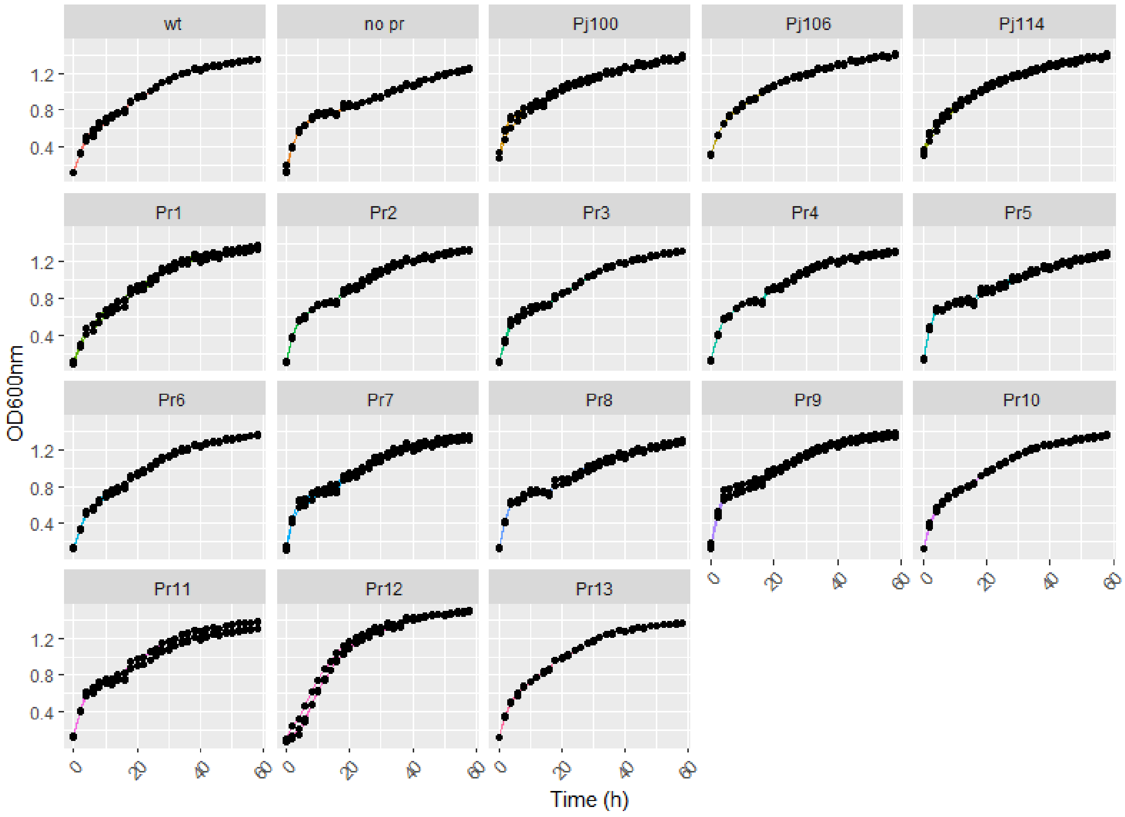

**Figure S6. Effect of plasmids in *C. taiwanensis* R1$^{T}$ fitness.** The effect on growth exerted by plasmids harboring the different promoter sequences and the different canonical promoters was evaluated. Strains were gown in LBKm at 25 °C for 16 h, diluted 20-fold in M9Km, and growth was evaluated at 25 °C every 2 h for a lapse of 60 h. Results are the mean of three technical replicates in the same experiment.

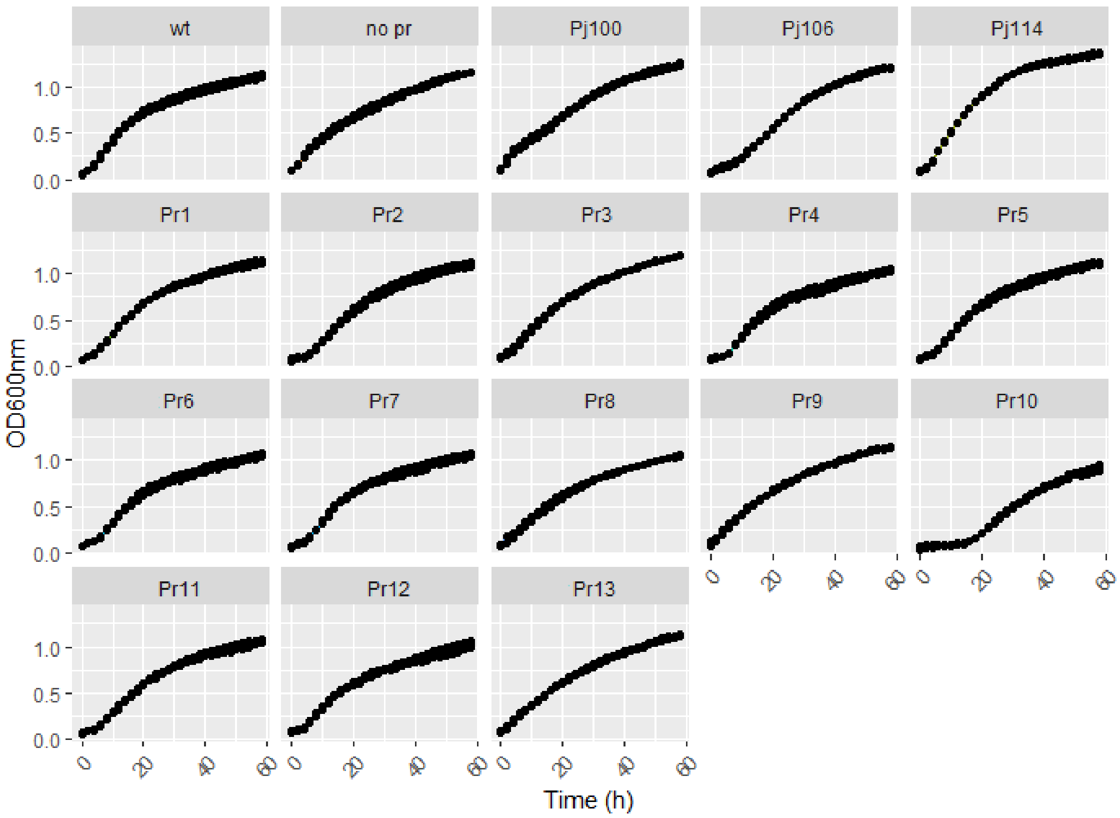

**Figure S7. Effect of plasmids in *P. phymatum* STM 815$^T$ fitness.** The effect on growth exerted by plasmids harboring the different promoter sequences and the different canonical promoters was evaluated. Strains were gown in LBKm at 25 °C for 16 h, diluted 20-fold in M9Km, and growth was evaluated at 25 °C every 2 h for a lapse of 60 h. Results are the mean of three technical replicates in the same experiment.

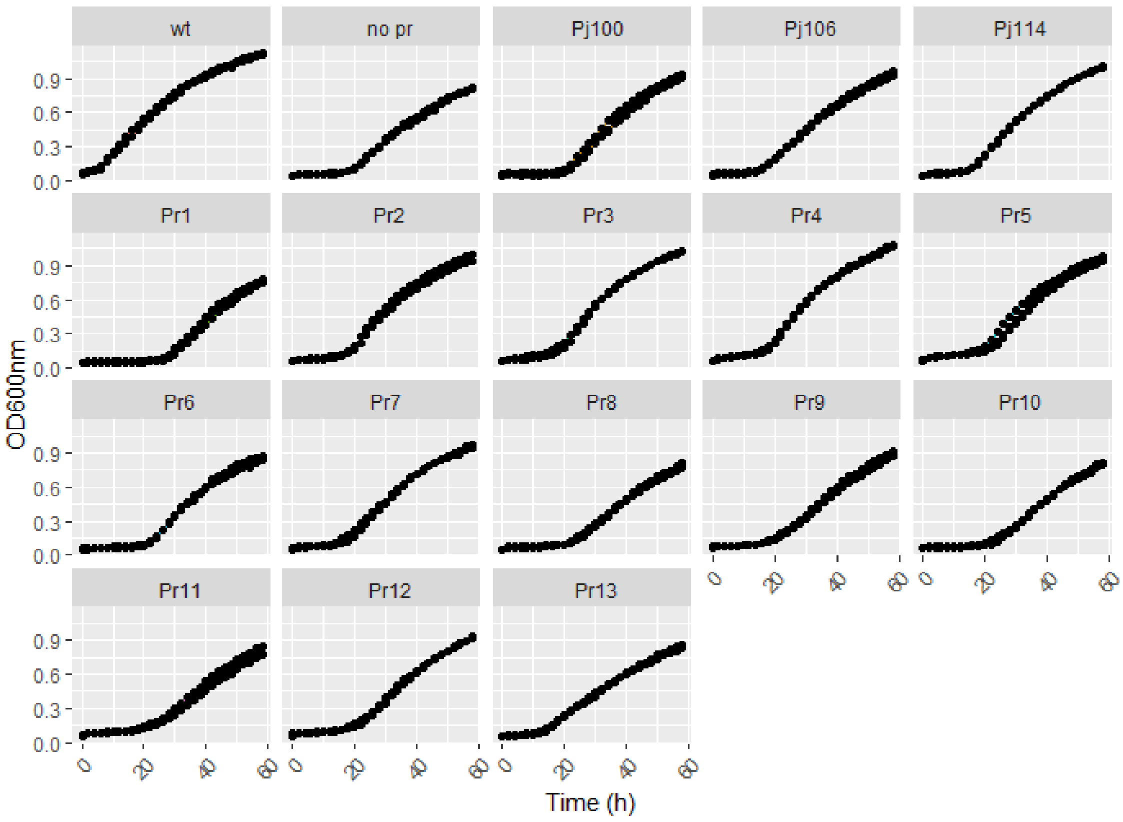

**Figure S8. Effect of plasmids in *E. meliloti* 1021 fitness.** The effect on growth exerted by plasmids harboring the different promoter sequences and the different canonical promoters was evaluated. Strains were gown in TYStrNm at 25 °C for 16 h, diluted 20-fold in M9SStrNm, and growth was evaluated at 25 °C every 2 h for a lapse of 60 h. Results are the mean of three technical replicates in the same experiment.

**Table S1.** Strains and plasmids

| Strain or plasmid | Relevant characteristics | Reference |
|---|---|---|
| *Pseudomonas putida* | | |
| KT2440 | Reference strain ($Km^S$) | (Bagdasarian et al. 1981) |
| KT2440 pSEVA231-*gfp* | KT2440 containing plasmid pSEVA231-*gfp* ($Km^R$) | (Amarelle et al. 2023) |
| KT2440 pSEVA231-*Pj100gfp* | KT2440 containing plasmid pSEVA231-*Pj100gfp* ($Km^R$) | (Amarelle et al. 2023) |
| KT2440 pSEVA231-*Pj106gfp* | KT2440 containing plasmid pSEVA231-*Pj106gfp* ($Km^R$) | (Amarelle et al. 2023) |
| KT2440 pSEVA231-*Pj114gfp* | KT2440 containing plasmid pSEVA231-*Pj114gfp* ($Km^R$) | (Amarelle et al. 2023) |
| KT2440 pSEVA231-*Pr(x)gfp* | KT2440 containing plasmids pSEVA231-*Pr(x)gfp,* where *x* refers to different metagenomic DNA sequences (1-13) exerting promoter activity ($Km^R$) | This work |
| *Pseudomonas* sp. | | |
| UYIF39 | Antarctic *Pseudomonas* isolated from a red mat in Fildes Peninsula of King George Island, South Shetland archipelago ($Km^S$) | (Ferrés et al. 2015) |
| UYIF39 pSEVA231-*gfp* | UYIF39 containing plasmid pSEVA231-*gfp* ($Km^R$) | (Amarelle et al. 2023) |
| UYIF39 pSEVA231-*Pj100gfp* | UYIF39 containing plasmid pSEVA231-*Pj100gfp* ($Km^R$) | (Amarelle et al. 2023) |
| UYIF39 pSEVA231-*Pj106gfp* | UYIF39 containing plasmid pSEVA231-*Pj106gfp* ($Km^R$) | (Amarelle et al. 2023) |
| UYIF39 pSEVA231-*Pj114gfp* | UYIF39 containing plasmid pSEVA231-*Pj114gfp* ($Km^R$) | (Amarelle et al. 2023) |
| UYIF39 pSEVA231- *Pr(x)gfp* | UYIF39 containing plasmids pSEVA231-*Pr(x)gfp,* where *x* refers to different metagenomic DNA sequences (1-13) ($Km^R$) | This work |
| *Escherichia coli* | | |
| DH5α | Φ80*lacZ*ΔM15 *recA1 endA1 gyrA96 thi-1 hsdR17*($r_K^- m_K^+$) *supE44 relA1 deoR* Δ(*lacZYA-argF*)*U169* ($Km^S$, $Str^S$) | (Hanahan 1983) |
| DH5α pRK2013 | DH5α containing helper plasmid pRK2013 ($Km^R$, $Str^S$) | (Hall et al. 2023) |
| DH5α pSEVA231-*gfp* | DH5α containing plasmid pSEVA231-*gfp* ($Km^R$) | This work |
| DH5α pSEVA231-*Pj100gfp* | DH5α containing plasmid pSEVA231-*Pj100gfp* ($Km^R$) | This work |
| DH5α pSEVA231-*Pj106gfp* | DH5α containing plasmid pSEVA231-*Pj106gfp* ($Km^R$) | This work |
| DH5α pSEVA231-*Pj114gfp* | DH5α containing plasmid pSEVA231-*Pj114gfp* ($Km^R$) | This work |
| DH5α pSEVA231- *Pr(x)gfp* | DH5α containing plasmids pSEVA231-*Pr(x)gfp,* where *x* refers to different metagenomic DNA sequences (1-13) ($Km^R$) | This work |
| *Cupriavidus taiwanensis* | | |
| $R1^T$ | Type strain of the species *C. taiwanensis* ($Km^S$) | (Vandamme 2004) |
| $R1^T$ pSEVA231-*gfp* | $R1^T$ containing plasmid pSEVA231-*gfp* ($Km^R$) | This work |
| $R1^T$ pSEVA231-*Pj100gfp* | $R1^T$ containing plasmid pSEVA231-*Pj100gfp* ($Km^R$) | This work |
| $R1^T$ pSEVA231-*Pj106gfp* | $R1^T$ containing plasmid pSEVA231-*Pj106gfp* ($Km^R$) | This work |

| | | |
|---|---|---|
| R1$^T$ pSEVA231-*Pj114gfp* | R1$^T$ containing plasmid pSEVA231-*Pj114gfp* (Km$^R$) | This work |
| R1$^T$ pSEVA231- *Pr(x)gfp* | R1$^T$ containing plasmids pSEVA231-*Pr(x)gfp,* where *x* refers to different metagenomic DNA sequences (1-13) (Km$^R$) | This work |
| *Paraburkholderia phymatum* | | |
| STM 815$^T$ | Type strain of the species *P. phymatum* (Km$^S$) | (Sawana et al. 2014) |
| STM 815$^T$ pSEVA231-*gfp* | STM 815$^T$ containing plasmid pSEVA231-*gfp* (Km$^R$) | This work |
| STM 815$^T$ pSEVA231-*Pj100gfp* | STM 815$^T$ containing plasmid pSEVA231-*Pj100gfp* (Km$^R$) | This work |
| STM 815$^T$ pSEVA231-*Pj106gfp* | STM 815$^T$ containing plasmid pSEVA231-*Pj106gfp* (Km$^R$) | This work |
| STM 815$^T$ pSEVA231-*Pj114gfp* | STM 815$^T$ containing plasmid pSEVA231-*Pj114gfp* (Km$^R$) | This work |
| STM 815$^T$ pSEVA231- *Pr(x)gfp* | STM 815$^T$ containing plasmids pSEVA231-*Pr(x)gfp,* where *x* refers to different metagenomic DNA sequences (1-13) (Km$^R$) | This work |
| *Ensifer meliloti* | | |
| 1021 | Streptomycin derivative of wild-type strain SU47 (Str$^R$) | (Meade et al. 1982) |
| 1021 pSEVA231-*gfp* | 1021 containing plasmid pSEVA231-*gfp* (Nm$^R$) | This work |
| 1021 pSEVA231-*Pj100gfp* | 1021 containing plasmid pSEVA231-*Pj100gfp* (Nm$^R$) | This work |
| 1021 pSEVA231-*Pj106gfp* | 1021 containing plasmid pSEVA231-*Pj106gfp* (Nm$^R$) | This work |
| 1021 pSEVA231-*Pj114gfp* | 1021 containing plasmid pSEVA231-*Pj114gfp* (Nm$^R$) | This work |
| 1021 pSEVA231- *Pr(x)gfp* | 1021 containing plasmids pSEVA231-*Pr(x)gfp,* where *x* refers to different metagenomic DNA sequences (1-13) (Nm$^R$) | This work |
| Plasmids | | |
| pRK2013 | ColE1 replicon with RK2 *tra* genes. Used for mobilizing incP and incQ plasmids (Km$^R$) | (Ditta et al. 1980) |
| pSEVA231 | Monofunction plasmid with pBBR1 replicon and MCS-default (Km$^R$) | (Silva-Rocha et al. 2013) |
| pSEVA231-*gfp* | pSEVA231 harboring *gfplva* from pMR1 cloned as a EcoRI/HindIII fragment (Km$^R$) | (Amarelle et al. 2023) |
| pSEVA231-*Pj100gfp* | pSEVA231 harboring the *Pj100-gfplva* construct from pMR1-*Pj100* cloned as a EcoRI/HindIII fragment (Km$^R$) | (Amarelle et al. 2023) |
| pSEVA231-*Pj106gfp* | pSEVA231 harboring the *Pj106-gfplva* construct from pMR1-*Pj106* cloned as a EcoRI/HindIII fragment (Km$^R$) | (Amarelle et al. 2023) |
| pSEVA231-*Pj114gfp* | pSEVA231 harboring the *Pj114-gfplva* construct from pMR1-*Pj114* cloned as a EcoRI/HindIII fragment (Km$^R$) | (Amarelle et al. 2023) |
| pSEVA231-*Pr1gfp* | pSEVA231-*gfp* harboring a 388 bp metagenomic DNA fragment cloned in BamHI restriction site (Km$^R$) | This work |
| pSEVA231-*Pr2gfp* | pSEVA231-*gfp* harboring a 306 bp metagenomic DNA fragment cloned in BamHI restriction site (Km$^R$) | This work |
| pSEVA231-*Pr3gfp* | pSEVA231-*gfp* harboring a 432 bp metagenomic DNA fragment cloned in BamHI restriction site (Km$^R$) | This work |
| pSEVA231-*Pr4gfp* | pSEVA231-*gfp* harboring a 695 bp metagenomic DNA fragment cloned in BamHI restriction site (Km$^R$) | This work |
| pSEVA231-*Pr5gfp* | pSEVA231-*gfp* harboring a 427 bp metagenomic DNA fragment cloned in BamHI restriction site (Km$^R$) | This work |

| | | |
|---|---|---|
| pSEVA231-*Pr6gfp* | pSEVA231-*gfp* harboring a 431 bp metagenomic DNA fragment cloned in BamHI restriction site ($Km^R$) | This work |
| pSEVA231-*Pr7gfp* | pSEVA231-*gfp* harboring a 242 bp metagenomic DNA fragment cloned in BamHI restriction site ($Km^R$) | This work |
| pSEVA231-*Pr8gfp* | pSEVA231-*gfp* harboring a 968 bp metagenomic DNA fragment cloned in BamHI restriction site ($Km^R$) | This work |
| pSEVA231-*Pr9gfp* | pSEVA231-*gfp* harboring a 481 bp metagenomic DNA fragment cloned in BamHI restriction site ($Km^R$) | This work |
| pSEVA231-*Pr10gfp* | pSEVA231-*gfp* harboring a 865 bp metagenomic DNA fragment cloned in BamHI restriction site ($Km^R$) | This work |
| pSEVA231-*Pr11gfp* | pSEVA231-*gfp* harboring a 441 bp metagenomic DNA fragment cloned in BamHI restriction site ($Km^R$) | This work |
| pSEVA231-*Pr12gfp* | pSEVA231-*gfp* harboring a 381 bp metagenomic DNA fragment cloned in BamHI restriction site ($Km^R$) | This work |
| pSEVA231-*Pr13gfp* | pSEVA231-*gfp* harboring a 369 bp metagenomic DNA fragment cloned in BamHI restriction site ($Km^R$) | This work |
| pSEVA231-*Pr3Agfp* | pSEVA231-*gfp* harboring a synthetized 52 bp sequence from *Pr3*, containing a promoter motif as suggested by *in silico* analysis with BDGP ($Km^R$) | This work |
| pSEVA231-*Pr5Agfp* | pSEVA231-*gfp* harboring a synthetized 52 bp sequence from *Pr5*, containing a promoter motif as suggested by *in silico* analysis with BDGP ($Km^R$) | This work |
| pSEVA231-*Pr5Bgfp* | pSEVA231-*gfp* harboring a synthetized 73 bp sequence from *Pr5*, containing a promoter motif as suggested by *in silico* analysis with BDGP ($Km^R$) | This work |
| pSEVA231-*Pr6Agfp* | pSEVA231-*gfp* harboring a synthetized 52 bp sequence from *Pr6*, containing a promoter motif as suggested by *in silico* analysis with BDGP ($Km^R$) | This work |
| pSEVA231-*Pr8Agfp* | pSEVA231-*gfp* harboring a synthetized 52 bp sequence from *Pr8*, containing a promoter motif as suggested by *in silico* analysis with BDGP ($Km^R$) | This work |
| pSEVA231-*Pr8Bgfp* | pSEVA231-*gfp* harboring a synthetized 52 bp sequence from *Pr8*, containing a promoter motif as suggested by *in silico* analysis with BDGP ($Km^R$) | This work |
| pSEVA231-*Pr8Cgfp* | pSEVA231-*gfp* harboring a synthetized 52 bp sequence from *Pr8*, containing a promoter motif as suggested by *in silico* analysis with BDGP ($Km^R$) | This work |
| pSEVA231-*Pr10Agfp* | pSEVA231-*gfp* harboring a synthetized 52 bp sequence from *Pr10*, containing a promoter motif as suggested by *in silico* analysis with BDGP ($Km^R$) | This work |
| pSEVA231-*Pr10Bgfp* | pSEVA231-*gfp* harboring a synthetized 52 bp sequence from *Pr10*, containing a promoter motif as suggested by *in silico* analysis with BDGP ($Km^R$) | This work |

**Table S2.** Primers used for the construction of the selected promoters

| **Fragment** | **Forward primer[a, c]**<br>**5'- 3'** | **Reverse primer[b,c]**<br>**5'- 3'** |
|---|---|---|
| ***Pr3A*** | **aattc**CGTTCGTTGCCCGCCCTCCTCAAA<br>ACGGTTATACTTATGCCAACCATGAG*g* | **gatcc**CTCATGGTTGGCATAAGTATAACC<br>GTTTTGAGGAGGGCGGGCAACGAACG*g* |
| ***Pr5A*** | **aattc**CTTCGATTGCCACGCCTTCTGCTGC<br>AGCCAATAATTCAAGGTCAGCAACA*g* | **gatcc**TGTTGCTGACCTTGAATTATTGGC<br>TGCAGCAGAAGGCGTGGCAATCGAAG*g* |
| ***Pr5B**** | **aattc**TCCCTCATGAATCTCACTTGACTG<br>CTTAGTCAGATAGAGCTAATATGACTA<br>TATAGTCAAGTAGGAATCTG*g* | **gatcc**CAGATTCCTACTTGACTATATAGT<br>CATATTAGCTCTATCTGACTAAGCAGT<br>CAAGTGAGATTCATGAGGGA*g* |
| ***Pr6A*** | **aattc**GGCTTGACATTGAACACTTAGTGG<br>ACCACTCTACTCCAAGTGTTCAATGC*g* | **gatcc**GCATTGAACACTTGGAGTAGAGTG<br>GTCCACTAAGTGTTCAATGTCAAGCC*g* |
| ***Pr8A*** | **aattc**AAGTTCCTGCCGGTCGATAGGGCG<br>GATGAGATAGTCGTTCACCCCGATGT*g* | **gatcc**ACATCGGGGTGAACGACTATCTCA<br>TCCGCCCTATCGACCGGCAGGAACTT*g* |
| ***Pr8B*** | **aattc**GCAGGTTTTGATGGAGCTGGAAGG<br>CCCGCTAAAAACGCACGCGGCCGAAG*g* | **gatcc**CTTCGGCCGCGTGCGTTTTTAGCG<br>GGCCTTCCAGCTCCATCAAAACCTGC*g* |
| ***Pr8C*** | **aattc**CTACTGTTGCGCAAATACCGCGTT<br>TTTGTTACAATGCCGAAACATTCTGT*g* | **gatcc**ACAGAATGTTTCGGCATTGTAACA<br>AAAACGCGGTATTTGCGCAACAGTAG*g* |
| ***Pr10A*** | **aattc**CCGGCTTCGACTGGTTGATGGGTC<br>TCGACTACCATTGGTTCTCCACGATG*g* | **gatcc**CATCGTGGAGAACCAATGGTAGTC<br>GAGACCCATCAACCAGTCGAAGCCGG*g* |
| ***Pr10B*** | **aattc**AGTCCATTGACATGTACGGCAATA<br>CAACGTACATTGTGAGAAGAAGCTGG*g* | **gatcc**CCAGCTTCTTCTCACAATGTACGT<br>TGTATTGCCGTACATGTCAATGGACT*g* |

[a] EcoRI restriction site is shown in bold lower case

[b] BamHI restriction site is shown in bold lower case

[c] The 3' guanine used to generate BamHI and EcoRI overhangs is shown in italics

[*] Selected sequence 5B is longer in order to cover the three promoter sequences predicted by BDGP in that particular region (Fig. S2)

**Table S3.** Promoter sequences identified *in silico* within the metagenomic DNA fragments cloned in the promoter-trap vector

| | Classification based on BacPP | | | | Classification based on SAPPHIRE | | | | Classification based BDGP | | | |
|---|---|---|---|---|---|---|---|---|---|---|---|---|
| **Metagenomic insert** | **Promoter** | **Promoter position** | **Promoter sequence** | **Sigma factor (accuracy %)** | **Promoter** | **Promoter position** | **Promoter sequence** | **p-value** | **Promoter** | **Promoter position** | **Promoter sequence** | **Score** |
| ***Pr1*** | **1** | 43-123 | CGATGAAATCTGGCT<br>GAAACTCTGGGGCA<br>ACCTCAGTTTCAATC<br>CGGTCAGCGTCCTGA<br>CCCATGGCACGCTTG<br>CGCAAT | S24 (15%),<br>S32 (19%),<br>S70 (54%) | **1** | 2-46 | GATCAACGCCGG<br>CATAAAAAGTCC<br>GGTGCGCGCCAA<br>TCTGCGCGA | 0.002801 | **1** | 52-97 | TCTGGCTGAAACTC<br>TGGGGCAACCTCA<br>GTTTCAATCCGGTC<br>AGCGTCCTG | 0,53 |
| | **2** | 72-152 | ACCTCAGTTTCAATC<br>CGGTCAGCGTCCTGA<br>CCCATGGCACGCTTG<br>CGCAATTGGCCACTG<br>ACGGCGGCACACGC<br>CGGGTC | S32 (52%) | **2** | 28-72 | TGCGCGCCAATCT<br>GCGCGATGAAAT<br>CTGGCTGAAACT<br>CTGGGGCA | 0.002559 | **2** | 302-347 | GGAACTCGACTCG<br>TTGCTGGGCGTTGT<br>CATAGAATTGGCA<br>CAGATGGTTG | 0,81 |

| | | | | | | | | | | | | |
|---|---|---|---|---|---|---|---|---|---|---|---|---|
| | **3** | 302-382 | GAACTCGACTCGTTGCTGGGCGTTGTCATAGAATTGGCACAGATGGTTGGAATTGCTACACCTGCCCTCAATTTGGTATA | S24 (10%), S38 (30%), S54 (61%), S70 (54%) | **3** | 38-82 | TCTGCGCGATGAAATCTGGCTGAAACTCTGGGGCAACCTCAGTTT | 0.001816 | **3** | 321-366 | GGCGTTGTCATAGAATTGGCACAGATGGTTGGAATTGCTACACCTGCCCT | 0.59 |
| | **4** | 303-383 | AACTCGACTCGTTGCTGGGCGTTGTCATAGAATTGGCACAGATGGTTGGAATTGCTACACCTGCCCTCAATTTGGTATAT | S38 (20%), S54 (54%), S70 (22%) | **4** | 48-92 | GAAATCTGGCTGAAACTCTGGGGCAACCTCAGTTTCAATCCGGTC | 0.004049 | | | | |
| | **5** | 305-385 | CTCGACTCGTTGCTGGGCGTTGTCATAGAATTGGCACAGATGGTTGGAATTGCTACACCTGCCCTCAATTTGGTATATGA | S24 (23%), S38 (30%), S70 (50%) | **5** | 78-122 | AGTTTCAATCCGGTCAGCGTCCTGACCCATGGCACGCTTGCGCAA | 0.003660 | | | | |
| | | | | | **6** | 128-172 | CACTGACGGCGGCACACGCCGGGTCATTCGTGCCATGATGCTGGA | 0.000280 | | | | |
| | | | | | **7** | 171-215 | GAGGCACAGGCGGTGGGCGAGGCGTTGGGCGTGAAATTTGCCGTG | 0.003012 | | | | |
| | | | | | **8** | 212-256 | CGTGGCGGTCGATGAGCGCATCGGCATGGCGGAAAAAGTCGGCAA | 0.004027 | | | | |

| | | | | | | | | | | | | |
|---|---|---|---|---|---|---|---|---|---|---|---|---|
| | | | | | **9** | 298-342 | CCACGGAACTCG ACTCGTTGCTGGG CGTTGTCATAGA ATTGGCAC | 0.000017 | | | | |
| | | | | | **10** | 327-371 | GTCATAGAATTG GCACAGATGGTT GGAATTGCTACA CCTGCCCTC | 0.003510 | | | | |
| ***Pr2*** | **1** | 211-291 | GGGTAGCCGAGCCC GACGCATCGCGTCTC AAAGCGAATTTTTGG CGGCCCTGTTGTCGA AATCGGTTGACGTGC GTAGGA | S32 (42%) | **1** | 67-111 | CGTCGACGCGCT CGTCACCAAGGT CGGGCAGCGAGA ATCGTCGGG | 0.004037 | **1** | 258-303 | GCCCTGTTGTCGAA ATCGGTTGACGTGC GTAGGATGTAGTC CTTGATCC | 0.92 |
| | | | | | **2** | 108-152 | CGGGCGCGATTG GTCGTCGGAGCA GTCATGGCTCATC CTGCGGGG | 0.000372 | | | | |
| | | | | | **3** | 255-299 | GCGGCCCTGTTGT CGAAATCGGTTG ACGTGCGTAGGA TGTAGTCC | 0.000002 | | | | |
| ***Pr3*** | **1** | 183-263 | GGCGGCCATAAACC GGACATTTCTACTGT GGGAGGAAGAGGAC ATTTCTAAAGTGGGT TGACAGGAAAGCAA CCGTTCGT | S32 (19%), S38 (20%), S70 (86%) | **1** | 20-64 | GCTCACTCACAA CGGTCGGGCACT CGGCTTTCACACA ATCGCGGC | 0.004196 | **1** | 257-302 | CGTTCGTTGCCCGC CCTCCTCAAAACG GTTATACTTATGCC AACCATGAG | 0.72 |

| | | | | | | | | |
|---|---|---|---|---|---|---|---|---|
| | **2** | 237-317 | GGGTTGACAGGAAAGCAACCGTTCGTTGCCCGCCCTCCTCAAAACGGTTATACTTATGCCAACCATGAGTCAGTTTCTTC | S38 (80%), S70 (72%) | **2** | 174-218 | GCCCCGACGGGGCGGCCATAAACCGGACATTTCTACTGTGGGAGG | 0.001092 |
| | **3** | 258-338 | TTCGTTGCCCGCCCTCCTCAAAACGGTTATACTTATGCCAACCATGAGTCAGTTTCTTCTTACTTTAAGCAGATTTACCG | S28 (49%), S38 (80%), S70 (45%) | **3** | 199-243 | GACATTTCTACTGTGGGAGGAAGAGGACATTTCTAAAGTGGGTTG | 0.000761 |
| | **4** | 259-339 | TCGTTGCCCGCCCTCCTCAAAACGGTTATACTTATGCCAACCATGAGTCAGTTTCTTCTTACTTTAAGCAGATTTACCGC | S38 (60%), S70 (77%) | **4** | 244-298 | AACCGTTCGTTGCCCGCCCTCCTCAAAACGGTTATACTTATGCCA | 0.000017 |
| | **5** | 270-350 | CCTCCTCAAAACGGTTATACTTATGCCAACCATGAGTCAGTTTCTTCTTACTTTAAGCAGATTTACCGCCCAGAAGCAGA | S24 (48%), S38 (60%), S70 (99%) | **5** | 266-310 | CCCGCCCTCCTCAAAACGGTTATACTTATGCCAACCATGAGTCAG | 0.002432 |
| | **6** | 279-359 | AACGGTTATACTTATGCCAACCATGAGTCAGTTTCTTCTTACTTTAAGCAGATTTACCGCCCAGAAGCAGACGGTCGACC | S24 (20%), S28 (45%), S32 (35%), S38 (55%), S70 (99%) | **6** | 287-331 | ATACTTATGCCAACCATGAGTCAGTTTCTTCTTACTTTAAGCAGA | 0.000790 |

| | | | | | | | | | |
|---|---|---|---|---|---|---|---|---|---|
| | **7** | 292-372 | ATGCCAACCATGAGTCAGTTTCTTCTTACTTTAAGCAGATTTACCGCCCAGAAGCAGACGGTCGACCATGAAACTGGAGG | S32 (61%), S38 (30%), S70 (77%) | **7** | 352-396 | GGTCGACCATGAAACTGGAGGGGGGCTTGCCCGTCTTCTGGGTCA | 0.003526 | |
| ***Pr4*** | **1** | 107-187 | TCGTAATCAACCGCGGCCGCCTCGAGTTCCGCCAGGATCGCTTCACCCTGTCGTCGCGCTACGCCGGGCGGCACTTCGGC | S32 (52%) | **1** | 55-99 | GCTCGGGATTCACCAGTTCAGGGCGCATTTCCTGTACCAGGTCGG | 0.002112 | **ND** |
| | **2** | 442-522 | CGAGGACTACGCGTTCGCTCGGCCGAGCCTCAACTCGCACAAGCGTCGCAAGCTCGAACTCGCGGCCGGTGATGCTTCGC | S54 (60%) | **2** | 78-122 | CGCATTTCCTGTACCAGGTCGGTGCCGTTCTCGTAATCAACCGCG | 0.003056 | |
| | | | | | **3** | 222-266 | CGCATCGCGATGTCCAGCAGGCCCTTATCGGTCAGGATCACCAAG | 0.000368 | |
| | | | | | **4** | 312-356 | GCGGCCGCTCGCACGCCAGGGCCGTTGCGGGCGTTCTATCGCCGG | 0.003473 | |
| | | | | | **5** | 521-565 | GCGGCGCGGGCCGGCCGCGGGGCCGTCGCGGGACTATCACGTCAA | 0.002329 | |

| | | | | | | | | | | | | |
|---|---|---|---|---|---|---|---|---|---|---|---|---|
| | | | | | **6** | 636-680 | TGGCAACCGAAG AGACCAACGGGC AACGGTTGACAA TCTCATCCG | 0.001357 | | | | |
| ***Pr5*** | **1** | 258-338 | ACCGTCATTGGGCCT CACTCCCTCATGAAT CTCACTTGACTGCTT AGTCAGATAGAGCT AATATGACTATATAG TCAAGT | S24 (15%), S70 (90%) | **1** | 2-46 | GATCTTCGATTGC CACGCCTTCTGCT GCAGCCAATAAT TCAAGGT | 0.001757 | **1** | 5-50 | CTTCGATTGCCACG CCTTCTGCTGCAGC CAATAATTCAAGG TCAGCAACA | 0.95 |
| | **2** | 268-348 | GGCCTCACTCCCTCA TGAATCTCACTTGAC TGCTTAGTCAGATAG AGCTAATATGACTAT ATAGTCAAGTAGGA ATCTGC | S32 (61%), S38 (99%), S70 (99%) | **2** | 71-115 | CCAGCGCCCGGA AGCCCGGCTCGC CTTTCTCGGTTAC ATGCAGGA | 0.004446 | **2** | 277-322 | TCCCTCATGAATCT CACTTGACTGCTTA GTCAGATAGAGCT AATATGACT | 0.69 |
| | **3** | 277-357 | CCCTCATGAATCTCA CTTGACTGCTTAGTC AGATAGAGCTAATAT GACTATATAGTCAAG TAGGAATCTGCCGAT GATGA | S24 (12%), S32 (42%), S38 (95%), S70 (72%) | **3** | 177-221 | TCCGAACTCGTTC AGGAACGCCCGC ATCAAGCCAAGA TCAGGCGC | 0.000789 | **3** | 288-333 | TCTCACTTGACTGC TTAGTCAGATAGA GCTAATATGACTAT ATAGTCAAG | 0.99 |
| | **4** | 279-359 | CTCATGAATCTCACT TGACTGCTTAGTCAG ATAGAGCTAATATGA CTATATAGTCAAGTA GGAATCTGCCGATGA TGAAA | S24 (41%), S32 (30%), S38 (45%), S70 (95%) | **4** | 273-317 | TCACTCCCTCATG AATCTCACTTGAC TGCTTAGTCAGAT AGAGCT | 0.004154 | **4** | 299-344 | TGCTTAGTCAGATA GAGCTAATATGAC TATATAGTCAAGTA GGAATCTGC | 0.76 |

| | | | | | | | | | | | | |
|---|---|---|---|---|---|---|---|---|---|---|---|---|
| | **5** | 292-372 | CTTGACTGCTTAGTCAGATAGAGCTAATATGACTATATAGTCAAGTAGGAATCTGCCGATGATGAAAACATTTGCCCGAC | S70 (90%) | **5** | 284-328 | TGAATCTCACTTGACTGCTTAGTCAGATAGAGCTAATATGACTAT | 0.000070 | | | | |
| | **6** | 293-373 | TTGACTGCTTAGTCAGATAGAGCTAATATGACTATATAGTCAAGTAGGAATCTGCCGATGATGAAAACATTTGCCCGACC | S24 (28%), S38 (10%), S70 (95%) | **6** | 294-338 | TTGACTGCTTAGTCAGATAGAGCTAATATGACTATATAGTCAAGT | 0.000826 | | | | |
| | **7** | 301-381 | TTAGTCAGATAGAGCTAATATGACTATATAGTCAAGTAGGAATCTGCCGATGATGAAAACATTTGCCCGACCCGCGCCAA | S24 (33%), S32 (54%), S38 (40%), S70 (90%) | **7** | 317-361 | TAATATGACTATATAGTCAAGTAGGAATCTGCCGATGATGAAAAC | 0.002131 | | | | |
| | **8** | 309-389 | ATAGAGCTAATATGACTATATAGTCAAGTAGGAATCTGCCGATGATGAAAACATTTGCCCGACCCGCGCCAAGGGCGGCC | S38 (90%), S70 (81%) | | | | | | | | |
| ***Pr6*** | **1** | 43-123 | CTCACCCAACAGCACAGGATTCGGCGTTCAGCAGGGGATTCTGAGCGCGACTAACCGTTCAATCCCGGGCTTGACATTGA | S24 (12%), S32 (66%) | **1** | 1-45 | GGATCTCGTGCGTAAACCGCCGAATCTCCTGCTGTAACCGGGCCT | 0.003683 | **1** | 111-156 | GGCTTGACATTGAACACTTAGTGGACCACTCTACTCCAAGTGTTCAATGC | 0.59 |

| | | | | | | | | |
|---|---|---|---|---|---|---|---|---|
| | **2** | 87-167 | GCGCGACTAACCGTTCAATCCCGGGCTTGACATTGAACACTTAGTGGACCACTCTACTCCAAGTGTTCAATGCGCCCAAT | S32 (71%), S38 (40%) | **2** | 25-69 | TCTCCTGCTGTAACCGGGCCTCACCCAACAGCACAGGATTCGGCG | 0.004236 |
| | **3** | 88-168 | CGCGACTAACCGTTCAATCCCGGGCTTGACATTGAACACTTAGTGGACCACTCTACTCCAAGTGTTCAATGCGCCCAATA | S24 (33%), S32 (42%), S38 (40%), S70 (54%) | **3** | 91-135 | CGACTAACCGTTCAATCCCGGGCTTGACATTGAACACTTAGTGGA | 0.003212 |
| | **4** | 89-169 | GCGACTAACCGTTCAATCCCGGGCTTGACATTGAACACTTAGTGGACCACTCTACTCCAAGTGTTCAATGCGCCCAATAG | S32 (52%), S38 (15%) | **4** | 104-148 | AATCCCGGGCTTGACATTGAACACTTAGTGGACCACTCTACTCCA | 0.002871 |
| | **5** | 91-171 | GACTAACCGTTCAATCCCGGGCTTGACATTGAACACTTAGTGGACCACTCTACTCCAAGTGTTCAATGCGCCCAATAGAC | S38 (50%) | **5** | 121-165 | TGAACACTTAGTGGACCACTCTACTCCAAGTGTTCAATGCGCCCA | 0.002502 |
| | **6** | 95-175 | AACCGTTCAATCCCGGGCTTGACATTGAACACTTAGTGGACCACTCTACTCCAAGTGTTCAATGCGCCCAATAGACCGAA | S24 (51%), S32 (35%), S70 (50%) | **6** | 250-294 | CGCGTCGTTCACCTACCGCGCATTCGTACCCGATCCTATCGCCGC | 0.001398 |

| | | | | |
|---|---|---|---|---|
| | **7** | 106-186 | CCCGGGCTTGACATTGAACACTTAGTGGACCACTCTACTCCAAGTGTTCAATGCGCCCAATAGACCGAACGGTTAGCAAT | S38 (10%), S70 (50%) |
| | **8** | 107-187 | CCGGGCTTGACATTGAACACTTAGTGGACCACTCTACTCCAAGTGTTCAATGCGCCCAATAGACCGAACGGTTAGCAATG | S70 (54%) |
| | **9** | 114-194 | TGACATTGAACACTTAGTGGACCACTCTACTCCAAGTGTTCAATGCGCCCAATAGACCGAACGGTTAGCAATGCCCCACG | S32 (50%), S70 (27%) |
| | **10** | 115-195 | GACATTGAACACTTAGTGGACCACTCTACTCCAAGTGTTCAATGCGCCCAATAGACCGAACGGTTAGCAATGCCCCACGA | S32 (52%) |
| | **11** | 118-198 | ATTGAACACTTAGTGGACCACTCTACTCCAAGTGTTCAATGCGCCCAATAGACCGAACGGTTAGCAATGCCCCACGAAGT | S38 (10%), S70 (54%) |

| | | | | | | | | | | | | |
|---|---|---|---|---|---|---|---|---|---|---|---|---|
| | **12** | 246-326 | TCGCGCGTCGTTCACCTACCGCGCATTCGTACCCGATCCTATCGCCGCCCTCGACCCCGACATCACCTTCGAGACGGCCG | S32 (59%) | | | | | | | | |
| ***Pr7*** | **1** | 69-149 | GCTCGCGGAACGATTCGAGTACCGCTTGCATCCTTGGACGAGTTTCGTCATCATCCCACTCTTCGCGCTCGCCAATGCAG | S32 (64%), S38 (35%) S70 (27%) | **1** | 22-66 | CGGTGCCGATGTGGCGGTCATGACCACCGTGGCAAAGTCGACGGT | 0.000951 | **1** | 89-134 | TACCGCTTGCATCCTTGGACGAGTTTCGTCATCATCCCACTCTTCGCGCT | 0.84 |
| | **2** | 76-156 | GAACGATTCGAGTACCGCTTGCATCCTTGGACGAGTTTCGTCATCATCCCACTCTTCGCGCTCGCCAATGCAGGCGTCAG | S32 (59%) | **2** | 66-110 | TGTCGCTCGCGGAACGATTCGAGTACCGCTTGCATCCTTGGACGA | 0.001449 | | | | |
| | **3** | 77-157 | AACGATTCGAGTACCGCTTGCATCCTTGGACGAGTTTCGTCATCATCCCACTCTTCGCGCTCGCCAATGCAGGCGTCAGA | S32 (54%), S38 (20%), S70 (40%) | **3** | 86-130 | GAGTACCGCTTGCATCCTTGGACGAGTTTCGTCATCATCCCACTC | 0.000385 | | | | |
| | **4** | 105-185 | GACGAGTTTCGTCATCATCCCACTCTTCGCGCTCGCCAATGCAGGCGTCAGACTCGATATGAACGCTTTCAGCGGGCCGG | S38 (75%), S70 (27%) | **4** | 122-166 | ATCCCACTCTTCGCGCTCGCCAATGCAGGCGTCAGACTCGATATG | 0.001335 | | | | |

| | | | | | | | | | | | | |
|---|---|---|---|---|---|---|---|---|---|---|---|---|
| ***Pr8*** | **1** | 675-755 | ATATGTGTCGATGTCATGAAACTTTACTCCCCTTGTCGGAATGGCCCCTACAACTGTGGCATTCCCTGCGATATCCCGAG | S32 (59%), S38 (85%), S70 (90%) | **1** | 22-66 | GGCCCAGATGGGTTTCCATGTAGCGCCGGTTGTAAAGGCCAGTGA | 0.004274 | **1** | 164-209 | AAGTTCCTGCCGGTCGATAGGGCGGATGAGATAGTCGTTCACCCCGATGT | 0.55 |
| | **2** | 779-859 | TTCCGCACGTCAAGACTTTAAGGAACTACTGTTGCGCAAATACCGCGTTTTTGTTACAATGCCGAAACATTCTGTGCGCG | S38 (60%), S70 (81%) | **2** | 128-172 | GCGGGCGCGGCGCACCTGGGTGTTCACCCTGGCTAGAAGTTCCTG | 0.003131 | **2** | 555-600 | GCAGGTTTTGATGGAGCTGGAAGGCCCGCTAAAAACGCACGCGGCCGAAG | 0.96 |
| | **3** | 785-865 | ACGTCAAGACTTTAAGGAACTACTGTTGCGCAAATACCGCGTTTTTGTTACAATGCCGAAACATTCTGTGCGCGTTTCAA | S24 (30%), S32 (35%), S38 (75%), S70 (99%) | **3** | 162-206 | AGAAGTTCCTGCCGGTCGATAGGGCGGATGAGATAGTCGTTCACC | 0.001437 | **3** | 805-850 | CTACTGTTGCGCAAATACCGCGTTTTTGTTACAATGCCGAAACATTCTGT | 0.77 |
| | **4** | 787-867 | GTCAAGACTTTAAGGAACTACTGTTGCGCAAATACCGCGTTTTTGTTACAATGCCGAAACATTCTGTGCGCGTTTCAAAG | S24 (46%), S32 (28%), S38 (65%), S54 (50%), S70 (100%) | **4** | 262-306 | GGATGGGAATTTGCCGGGTCTGCTCCAGCGATTTCAGTTGCGAGC | 0.003785 | | | | |
| | **5** | 794-874 | CTTTAAGGAACTACTGTTGCGCAAATACCGCGTTTTTGTTACAATGCCGAAACATTCTGTGCGCGTTTCAAAGTCACTCA | S24 (30%), S32 (21%), S38 (75%), S70 (22%) | **5** | 552-596 | GGCGCAGGTTTTGATGGAGCTGGAAGGCCCGCTAAAAACGCACGC | 0.000663 | | | | |

| | | | | | | | | |
|---|---|---|---|---|---|---|---|---|
| | **6** | 796-876 | TTAAGGAACTACTGTTGCGCAAATACCGCGTTTTTGTTACAATGCCGAAACATTCTGTGCGCGTTTCAAAGTCACTCAGT | S24 (15%), S32 (52%), S38 (75%), S70 (90%) | **6** | 595-639 | GCGGCCGAAGCCGCCGACGGCATGGCAACGGCAATCATACCGCCC | 0.002685 |
| | **7** | 828-908 | TTTGTTACAATGCCGAAACATTCTGTGCGCGTTTCAAAGTCACTCAGTCCAGTTGTTCTTCAGGACTGATGCCGTGGATT | S38 (75%), S70 (81%) | **7** | 641-685 | AGCCCAAAGCGGCGCCACTCAGCAGCCGTGCGCGAATATGTGTCG | 0.000850 |
| | | | | | **8** | 662-706 | GCAGCCGTGCGCGAATATGTGTCGATGTCATGAAACTTTACTCCC | 0.002018 |
| | | | | | **9** | 692-736 | TGAAACTTTACTCCCCTTGTCGGAATGGCCCCTACAACTGTGGCA | 0.002659 |
| | | | | | **10** | 735-779 | CATTCCCTGCGATATCCCGAGCGGAACGGTGTATCGAATGCAGAG | 0.001112 |
| | | | | | **11** | 774-818 | GCAGAGTTCCGCACGTCAAGACTTTAAGGAACTACTGTTGCGCAA | 0.002868 |
| | | | | | **12** | 802-846 | GAACTACTGTTGCGCAAATACCGCGTTTTTGTTACAATGCCGAAA | 0.000000 |

| | | | | | | | | | | | | |
|---|---|---|---|---|---|---|---|---|---|---|---|---|
| ***Pr9*** | **1** | 351-431 | CATCGGATTGTGCGGGAGGAACCCTACACATCTGTGTTGTTCTGGCCCATCATTTCATGTATGGTCGACACAACTTCGGA | S24 (10%), S32 (76%), S38 (40%), S70 (40%) | **1** | 12-56 | GTCGTCGTCGTGTTGGTCGATGACGGCTCGGCCACGCTCGTTCTC | 0.004220 | **1** | 382-427 | TCTGTGTTGTTCTGGCCCATCATTTCATGTATGGTCGACACAACTTCGGA | 0.64 |
| | **2** | 362-442 | GCGGGAGGAACCCTACACATCTGTGTTGTTCTGGCCCATCATTTCATGTATGGTCGACACAACTTCGGACGAACTAGCAC | S32 (16%), S38 (30%), S70 (86%) | **2** | 140-184 | GAGCAGCATCAGCCGTCGCAGCGCGGCAGTAGGAATCATCACCAA | 0.004428 | | | | |
| | **3** | 374-454 | CTACACATCTGTGTTGTTCTGGCCCATCATTTCATGTATGGTCGACACAACTTCGGACGAACTAGCACGAAAGGCCTAGA | S32 (78%), S70 (27%) | **3** | 161-205 | CGCGGCAGTAGGAATCATCACCAACCTGCTGGGCCTAATCGAGGG | 0.001176 | | | | |
| | | | | | **4** | 366-410 | GGAGGAACCCTACACATCTGTGTTGTTCTGGCCCATCATTTCATG | 0.000566 | | | | |
| | | | | | **5** | 379-423 | ACATCTGTGTTGTTCTGGCCCATCATTTCATGTATGGTCGACACA | 0.000359 | | | | |

| | | | | | | | | | | | |
|---|---|---|---|---|---|---|---|---|---|---|---|
| ***Pr10*** | **1** | 515-595 | GTGCGTTTTTCTTAGCAGCAAAACAGTCCATTGACATGTACGGCAATACAACGTACATTGTGAGAAGAAGCTGGTGGGTC | S24 (12%), S32 (19%), S38 (30%), S70 (81%) | **1** | 36-80 | CGCACGGCTTCTTCTACGTGCGGATGGTCGTTTACTTTGTGTTGC | 0.000678 | **1** | 213-258 | CCGGCTTCGACTGGTTGATGGGTCTCGACTACCATTGGTTCTCCACGATG | 0.66 |
| | **2** | 520-600 | TTTTTCTTAGCAGCAAAACAGTCCATTGACATGTACGGCAATACAACGTACATTGTGAGAAGAAGCTGGTGGGTCACGAT | S24 (12%), S32 (30%), S38 (85%), S70 (95%) | **2** | 209-253 | TTTGCCGGCTTCGACTGGTTGATGGGTCTCGACTACCATTGGTTC | 0.000230 | **2** | 540-585 | AGTCCATTGACATGTACGGCAATACAACGTACATTGTGAGAAGAAGCTGG | 0.96 |
| | | | | | **3** | 223-267 | CTGGTTGATGGGTCTCGACTACCATTGGTTCTCCACGATGTGGGG | 0.003651 | | | | |
| | | | | | **4** | 239-283 | GACTACCATTGGTTCTCCACGATGTGGGGTGTTTATCTCTTCGCC | 0.004166 | | | | |
| | | | | | **5** | 346-390 | CTATCTCAAGGTCGTGAACACCGAGCATTACCACATCATGGGTAA | 0.003569 | | | | |
| | | | | | **6** | 355-399 | GGTCGTGAACACCGAGCATTACCACATCATGGGTAAGTTCATGCT | 0.003728 | | | | |
| | | | | | **7** | 375-419 | ACCACATCATGGGTAAGTTCATGCTGGCCTTCACCATTTTCTGGG | 0.002202 | | | | |

| | | | | | | | | |
|---|---|---|---|---|---|---|---|---|
| | | | | | **8** | 414-458 | TCTGGGCCTACATCGGCTTCTGCCAGTACATGCTCATCTGGTATG | 0.002212 |
| | | | | | **9** | 428-472 | GGCTTCTGCCAGTACATGCTCATCTGGTATGCGAACATTCCGGAG | 0.002587 |
| | | | | | **10** | 448-492 | CATCTGGTATGCGAACATTCCGGAGGAAACGATCACCCTGTGGTC | 0.002800 |
| | | | | | **11** | 492-536 | CACTGAACAGTAGGGGCCGCCGGAGTGCGTTTTTCTTAGCAGCAA | 0.003510 |
| | | | | | **12** | 538-582 | ACAGTCCATTGACATGTACGGCAATACAACGTACATTGTGAGAAG | 0.000155 |
| | | | | | **13** | 547-591 | TGACATGTACGGCAATACAACGTACATTGTGAGAAGAAGCTGGTG | 0.004516 |
| | | | | | **14** | 562-606 | TACAACGTACATTGTGAGAAGAAGCTGGTGGGTCACGATGAAACT | 0.000791 |

| | | | | | | | | | | | | |
|---|---|---|---|---|---|---|---|---|---|---|---|---|
| | | | | | **15** | 693-737 | GCTCGGTCAGCG ATTTCGTCGTGGC CGCCATACAGGA TGCCGCGC | 0.004460 | | | | |
| | | | | | **16** | 757-801 | GGAAGTCATCCG GCTCTCACGCGA GGGGCAGGAAAA ATTCGTGTC | 0.003723 | | | | |
| | | | | | **17** | 818-862 | CCGCCTTCGTTCG CGGCTCTTGCGCG CGCAATTAAACG TCACAGA | 0.002826 | | | | |
| ***Pr11*** | **1** | 98-178 | TTCAAACTCGCCGTC ATCGACGACGTCAAC AACCGCACCAACCC GCCCGACCTCACCTT CGCCGACTCATGGCG TCCGGT | S32 (54%) | **1** | 24-68 | AAGCGGGTCTGG ATTCCCTTGATGG GACTGGTCAGCTT CGTATCG | 0.002171 | **1** | 375-420 | TAGAGCTTGACGTC AAACGCCAACTGC TCAATCATCCTATC CACTACGCC | 0.75 |
| | **2** | 212-292 | AACATCATGGCCCGC GAGGAGGGCCACAA AGCCGTGAAGCTCCT CATGCTCGTGGTGGA TAACGACAACGCGG TGGTGAT | S54 (50%) | **2** | 69-113 | AGCGCGGCGCGA CCGTTGGGCGCC AGCGATTTCAAA CTCGCCGTC | 0.002124 | | | | |
| | | | | | **3** | 182-226 | CCGCGTGCGGGA GCGCAACGGAGA GTCCGTGAACAT CATGGCCCG | 0.001110 | | | | |
| | | | | | **4** | 373-417 | GTTAGAGCTTGA CGTCAAACGCCA ACTGCTCAATCAT CCTATCCA | 0.001176 | | | | |

| | | | | | | | | | | | | |
|---|---|---|---|---|---|---|---|---|---|---|---|---|
| ***Pr12*** | **1** | 7-87 | CCAGCCAAATGTTCTCAGCGTCCATGCGGTGCTGGCCGGTTTCTTCGCCATAAAAACCATAGCTCGGAAAGCGCTGCGCC | S24 (23%), S32 (57%), S70 (22%) | **1** | 25-69 | GCGTCCATGCGGTGCTGGCCGGTTTCTTCGCCATAAAAACCATAG | 0.003475 | **1** | 261-306 | TCTGCCTTGACAGCCCGCGGCGCGCCATTGAACATGCGCCGGTCGAGAGG | 0.66 |
| | **2** | 67-147 | AGCTCGGAAAGCGCTGCGCCAGCACTTCGTGAATGGCCTCCTCGGCGCGAACATCCGCTTCAGTGACGGGTGAACGGTCG | S32 (50%) | **2** | 223-267 | GCCGCCTGGAGTTCGGGGGAGGGCATGGCGCGGACAATTCTGCCT | 0.001682 | | | | |
| | **3** | 137-217 | TGAACGGTCGGATTTGATCTGCACTTTGAGGTTGCGCTGATACAGCCCGCGAATCACTTCGGCAGCGGCGGCGGCGGCGT | S24 (10%), S32 (66%) | **3** | 258-302 | AATTCTGCCTTGACAGCCCGCGGCGCGCCATTGAACATGCGCCGG | 0.001099 | | | | |
| ***Pr13*** | **1** | 73-153 | CCGAGTACTCGTCGAACTGGATCGGGTCGTAGTTCCACATGCTCATGGCGAAGGGCCCGTCGGCAAGGATGCCAGGCGGC | S28 (56%) | **1** | 106-150 | TTCCACATGCTCATGGCGAAGGGCCCGTCGGCAAGGATGCCAGGC | 0.002218 | | | | |
| | | | | | **2** | 272-316 | GCGCGAAGCGCTGGGTGTCGAACGTGATGCGCTCCATCTGGCGGA | 0.004027 | | | | |

ND, no promoter detected

**Table S4.** DNA sequences of the active metagenomic fragments showing promoters predicted by BDGP and their features

| **Metagenomic insert** | **5'- Sequence -3'**[ab] | **Identified promoters by BDGP** | **-35/-10 elements** [a] |
|---|---|---|---|
| *Pr1* (388 bp) | GGATCAACGCCGGCATAAAAAGTCCGGTGCGCG<br>CCAATCTGCGCGATGAAATCTGG**CTGAAA**CTCT<br>GGGGCAACCTCAGT**TTCAAT**CCGGTCAGCGTCC<br>TGACCCATGGCACGCTTGCGCAATTGGCCACTG<br>ACGGCGGCACACGCCGGGTCATTCGTGCCATGA<br>TGCTGGAGGCACAGGCGGTGGGCGAGGCGTTGG<br>GCGTGAAATTTGCCGTGGCGGTCGATGAGCGCA<br>TCGGCATGGCGGAAAAAGTCGGCAATCACCGCA<br>CGTCCATGCTGCAGGACGTGGAAGCGGGACGGC<br>CCACGGAAC**TCGACT**CGTTGCTGGGCG**TTGTCA**<br>**TAGAAT**TGGCACAGATGGT**TGGAAT**TGCTACAC<br>CTGCCCTCAATTTGGTATATGATCC | *Pr1A* (50 bp) | CTGAAA-$N_{18}$-TTCAAT |
| | | *Pr1B* (50 bp) | TCGACT-$N_{18}$-TAGAAT<br>TTGTCA-$N_{19}$-TGGAAT |
| *Pr2* (306 bp) | GGATCGCAGCCGCGCTGGCCTCGCCCTCGGGAT<br>GCTCCGGGCCGCTCATGAACGCCGTGATCGCCT<br>CGTCGACGCGCTCGTCACCAAGGTCGGGCAGCG<br>AGAATCGTCGGGCGCGATTGGTCGTCGGAGCAG<br>TCATGGCTCATCCTGCGGGGTGTGCGAGTCCGG<br>CGATGGTGCGGGCCGAAACGGCCTGCGTGTGCC<br>GGGTGGAGTTCAAGGGTAGCCGAGCCCGACGCA<br>TCGCGTCTCAAAGCGAATTTTTGGCGGCCCTG**TT**<br>**GTCG**AAATCGGTTGACGTGCG**TAGGAT**GTAGTC<br>CTTGATCC | *Pr2A* (50 bp) | TTGTCG-$N_{17}$-TAGGAT |
| *Pr3* (432 bp) | GGATCGATGGGACGATGCGGCTCACTCACAACG<br>GTCGGGCACTCGGCTTTCACACAATCGCGGCTC<br>GTCCCGTGAAGGCGACGGTCACCCCGGTCCACA<br>CGCCTCAGTGCCCGGTCACGCCGAGGCCGGACC<br>ATCCATGGCGCACACGCCTGCGGCCGGAGCGAA<br>TAACACAGGCCCCGACGGGGCGGCCATAAACCG<br>GACATTTCTACTGTGGGAGGAAGAGGACATTTC<br>TAAAGTGGGTTGACAGGAAAGCAACCGTTCG**TT**<br>**GCCC**GCCCTCCTCAAAACGGT**TATACT**TATGCC<br>AACCATGAGTCAGTTTCTTCTTACTTTAAGCAGA<br>TTTACCGCCCAGAAGCAGACGGTCGACCATGAA<br>ACTGGAGGGGGGCTTGCCCGTCTTCTGGGTCAG<br>ATCGGTCTGGTTGGGAAATGTATTGCGCAGGAT<br>CC | *Pr3A* (50 bp) | TTGCCC-$N_{17}$-TATACT |
| *Pr5* (427 bp) | GGATCTTCGA**TTGCCA**CGCCTTCTGCTGCAGCC<br>**AATAAT**TCAAGGTCAGCAACACTTTCCGCACGC<br>AACCCCAGCGCCCGGAAGCCCGGCTCGCCTTTC<br>TCGGTTACATGCAGGAACGGGGCACCGCCCAAT<br>CCGTGTGCGAACAGGCGTTCGCTTGTGCTGTTGA<br>CGCATTCAAGTCCGAACTCGTTCAGGAACGCCC<br>GCATCAAGCCAAGATCAGGCGCGCCAAAGCGTA<br>CATGCGCGATGTCTTCGATCCTGATCACCGTCAT<br>TGGGCCTCACTCCCTCATGAATCTCAC**TTGACT**<br>GC**TTAGTC**AGATAGAGC**TAATAT**GAC**TATATA**<br>GTCAAGTAGGAATCTGCCGATGATGAAAACATT<br>TGCCCGACCCGCGCCAAGGGCGGCCCGGACGAG<br>GCAGGCATTGCTCGCTGCAGGTTTCGATCC | *Pr5A* (50 bp) | TTGCCA-$N_{17}$-AATAAT |
| | | *Pr5B* (71 bp) | TTGACT-$N_{17}$-TAATAT<br>TTAGTC-$N_{18}$-TATATA |
| *Pr6* (431 bp) | GGATCTCGTGCGTAAACCGCCGAATCTCCTGCT<br>GTAACCGGGCCTCACCCAACAGCACAGGATTCG<br>GCGTTCAGCAGGGGATTCTGAGCGCGACTAACC<br>GTTCAATCCCGGGCTTGACA**TTGAAC**ACTTAGT<br>GGACCACTC**TACTCC**AAGTGTTCAATGCGCCCA<br>ATAGACCGAACGGTTAGCAATGCCCCACGAAGT | *Pr6A* (50 bp) | TTGAAC-$N_{16}$-TACTCC |

| | | | |
|---|---|---|---|
| | CGAGCGTCGATGGCAGGGCGACCCGGGCGCACC<br>AGGCGGACGTCGCGCTCGCGCGTCGTTCACCTA<br>CCGCGCATTCGTACCCGATCCTATCGCCGCCCTC<br>GACCCCGACATCACCTTCGAGACGGCCGAGTCG<br>ATTGCCCGGGCAGAGGCGGCAATCACCGACCTC<br>AACACCGCCGATTCGGTCGCTGGGCTCGAGGCC<br>ATCGGACCGCTGCTGCTGCGCTCGGAGGCGATC<br>C | | |
| *Pr7* (242 bp) | GGATCTTGAAGACGAACCAACCGGTGCCGATGT<br>GGCGGTCATGACCACCGTGGCAAAGTCGACGGT<br>GTCGCTCGCGGAACGATTCGAGTACCGCTTGCA<br>TCCTTGGACGAGTTTCGTCATCATCCCACTCTT<br>CGCGCTCGCCAATGCAGGCGTCAGACTCGATAT<br>GAACGCTTTCAGCGGGCCGGGAACGGCAGCCGT<br>TGGGGGCGGCATTGCCCTCGGACTCGTAGCCGG<br>AAAGACGATCC | *Pr7A* (50 bp) | **TTG**CAT-$N_{17}$-CA**T**CAT<br>**TTG**GAC-$N_{17}$-CA**CT**CT |
| *Pr8* (968 bp) | GGATCGAATGCTCAACCAGATGGCCCAGATGGG<br>TTTCCATGTAGCGCCGGTTGTAAAGGCCAGTGA<br>GCGCATCGGTGACCGCCATTTCAATCGAGGTCT<br>GCACGCTGGCGCGCAGCTGCTCGGCATAGCGGG<br>CGCGGCGCACCTGGGTGTTCACCCTGGCTAGAA<br>GTTCCTGCCGGTCGATAGGGCGGATGAGATAG<br>TCGTTCACCCCGATGTCAAGCGCCCGCATGAGG<br>CGTTGCTCGTCATCGGGCGCAACAACGATGAGG<br>ATGGGAATTTGCCGGGTCTGCTCCAGCGATTTCA<br>GTTGCGAGCACAGGCGCAGGCCATCGAACTGCT<br>CGGCATCCAGATTGACGATGACCAGTTCAAAGG<br>GGGTCGAGCTTTCCGTAATCGACGGAATGACAT<br>CCTGCGGATCAATGACCGCAGAAACCTTGTGCT<br>TGCCGGCAAGTGCCACGCGCACCCGGTCGATGA<br>ACGCCACGCGGCTGTCAACCACCAGGATATTGC<br>CATGGCGCGTGTCGATCGACGGGGCCCAGACCG<br>TTCGTGTCGGTCGTGGTCGTCGTGGCGCAGGTTT<br>TGATGGAGCTGGAAGGCCCGCTAAAAACGCAC<br>GCGGCCGAAGCCGCCGACGGCATGGCAACGGC<br>AATCATACCGCCCAAGCCCAAAGCGGCGCCACT<br>CAGCAGCCGTGCGCGAATATGTGTCGATGTCAT<br>GAAACTTTACTCCCCTTGTCGGAATGGCCCCTAC<br>AACTGTGGCATTCCCTGCGATATCCCGAGCGGA<br>ACGGTGTATCGAATGCAGAGTTCCGCACGTCAA<br>GACTTTAAGGAACTACTGTTGCGCAAATACCGC<br>GTTTTTGTTACAATGCCGAAACATTCTGTGCGCG<br>TTTCAAAGTCACTCAGTCCAGTTGTTCTTCAGGA<br>CTGATGCCGTGGATTTCGGCCTTCGCTATCCAGC<br>CGTTGCGATTGCCAATCGAGAACTTGCACCAAT<br>TTGGATCC | *Pr8A* (50 bp) | **TT**CCTG-$N_{17}$-**T**GAGA**T** |
| | | *Pr8B* (50 bp) | **TTGA**TG-$N_{16}$-**TA**AAAA |
| | | *Pr8C* (50 bp) | **TTG**CGC-$N_{17}$-**TA**CAA**T** |
| *Pr9* (481 bp) | GGATCTTCATCGTCGTCGTCGTGTTGGTCGATGA<br>CGGCTCGGCCACGCTCGTTCTCGGCGTCGACGA<br>CCTTGGTGACCGCGTCGTGGCTGTCCTTGTACCG<br>ACTCACACGGCACCGCCCACCTCGACGGGCTGC<br>TCATCGAGCAGCATCAGCCGTCGCAGCGCGGCA<br>GTAGGAATCATCACCAACCTGCTGGGCCTAATC<br>GAGGGCACGTCGCCGCGCGCTACGTCGTACGTG<br>GCAGAGCGGCCGAGACCGAGGAAGTGGCGGGC<br>TATCTCGGGCTACAGAACGGCATTCACCTCGGG<br>ATCCGGCAGGGTTGAGGACTCAGGCAATGTGCC<br>TCCTTCGTGATGGACAGACCCATCGGATTGTGC<br>GGGAGGAACCCTACACATCTGTGTTGTTCTGGC<br>CCATCATTTCATGTATGGTCGACACAACTTCGG<br>ACGAACTAGCACGAAAGGCCTAGAGGTTGATGG<br>CAGCCACCAAGAAGATCC | *Pr9A* (50 bp) | **TTG**TTC-$N_{17}$-**TA**TGG**T** |

| | | | |
|---|---|---|---|
| *Pr10* (865 bp) | GGATCCTTTGCTGGATGCCAAGCAACCCTATCTC<br>TCGCACGGCTTCTTCTACGTGCGGATGGTCGTTT<br>ACTTTGTGTTGCTCGGTTGGATCGGCTGGAGCCA<br>GTGGAAGCTATCCACTTCCCAAGACAAGGATGG<br>CGCGGCGAAGCACACCCACTTGATGCGCAAGTT<br>CGGCATTGGCGGCATCCCGGCGCTCGCACTGTG<br>CATCACCTTTGCCGGCT**TCGACT**GGTTGATGGG<br>TCTCGAC**TACCAT**TGGTTCTCCACGATGTGGGG<br>TGTTTATCTCTTCGCCGGCGCGGCGGGCAGTTCC<br>ATGTCTCTCCTCGTGCTCGTCGTGACGTGGTTGA<br>AATCGAAGGGCTATCTCAAGGTCGTGAACACCG<br>AGCATTACCACATCATGGGTAAGTTCATGCTGG<br>CCTTCACCATTTTCTGGGCCTACATCGGCTTCTG<br>CCAGTACATGCTCATCTGGTATGCGAACATTCCG<br>GAGGAAACGATCACCCTGTGGTCACTGAACAGT<br>AGGGGCCGCCGGAGTGCGTTTTTCTTAGCAGCA<br>AAACAGTCCA**TTGACA**TGTACGGCAATACAACG<br>**TACATT**GTGAGAAGAAGCTGGTGGGTCACGATG<br>AAACTACAAGAACCACGCACCGCTCGTTTGGAA<br>GCCCGCATCGCGCCCGAAACCCTCGCCATCGTC<br>CGGCGCGCGGCCGAGATTCAGGGCCGCTCGGTC<br>AGCGATTTCGTCGTGGCCGCCATACAGGATGCC<br>GCGCGACGGACGGTAGCCGAAATGGAAGTCATC<br>CGGCTCTCACGCGAGGGGCAGGAAAAATTCGTG<br>TCTTTGTTGATGAATCCTCCGCCTTCGTTCGCGG<br>CTCTTGCGCGCGCAATTAAACGTCACAGATCC | *Pr10A* (50 bp) | **TCGACT**-N$_{17}$-**TACCAT** |
| | | *Pr10B* (50 bp) | **TTGACA**-N$_{17}$-**TACATT** |
| *Pr11* (441 bp) | GGATCGAATCACACTACCAGAAGAAGCGGGTCT<br>GGATTCCCTTGATGGGACTGGTCAGCTTCGTATC<br>GAGCGCGGCGCGACCGTTGGGCGCCAGCGATTT<br>CAAACTCGCCGTCATCGACGACGTCAACAACCG<br>CACCAACCCGCCCGACCTCACCTTCGCCGACTC<br>ATGGCGTCCGGTGATCCGCGTGCGGGAGCGCAA<br>CGGAGAGTCCGTGAACATCATGGCCCGCGAGGA<br>GGGCCACAAAGCCGTGAAGCTCCTCATGCTCGT<br>GGTGGATAACGACAACGCGGTGGTGATGCAGAT<br>GCGCATGGCGCCGACCCGCTTCCTTGAGTTTGTG<br>GCCGAGCAAGCGCGCAACAAGCACCGCCGCGAT<br>GATGACCGTTAGAGC**TTGACG**TCAAACGCCAAC<br>TGCTC**AATCAT**CCTATCCACTACGCCGCCAGCG<br>ACAGCGATCC | *Pr11A* (50 bp) | **TTGACG**-N$_{17}$-**AATCAT** |
| *Pr12* (381 bp) | GGATCGACCAGCCAAATGTTCTCAGCGTCCATG<br>CGGTGCTGGCCGGTTTCTTCGCCATAAAAACCAT<br>AGCTCGGAAAGCGCTGCGCCAGCACTTCGTGAA<br>TGGCCTCCTCGGCGCGAACATCCGCTTCAGTGA<br>CGGGTGAACGGTCGGATTTGATCTGCACTTTGA<br>GGTTGCGCTGATACAGCCCGCGAATCACTTCGG<br>CAGCGGCGGCGGCGGCGTCGAGCGCCGCCTGGA<br>GTTCGGGGGAGGGCATGGCGCGGACAATTCTGC<br>C**TTGACA**GCCCGCGGCGCGCCATT**GAACAT**GCG<br>CCGGTCGAGAGGTGTTCGGGAAGTCGGTGAAGC<br>ACTTGGTGCCATTCCGACGCTGCCCCCGCAACG<br>GTAAGCAGGTTTGATCC | *Pr12A* (50 bp) | **TTGACA**-N$_{17}$-**GAACAT** |

[a] Elements -35 are depicted in red bold letters; elements -10 are depicted in blue bold letters
[b] Promoters sequences detected by BDGP are highlighted in gray

**Table S5.** Similarity in the amino acid sequences of the transcription factor $\sigma^{70}$ or RpoD in the genomes of the strains used

| | *P. putida* KT2440 | *Pseudomonas* sp. UYIF39 | *E. coli* DH5α | *C. taiwanensis* R1$^{T}$ [a] | *P. phymatum* STM 815$^{T}$ [a] | *E. meliloti* 1021 |
|---|---|---|---|---|---|---|
| *P. putida* KT2440 GCF_900167985.1 | 1 | 87.34 | 67.10 | 47.90-62.37 | 55.36-54.85 | 48.72 |

[a] For the *C. taiwanensis* R1$^{T}$ and *P. phymatum* STM 815$^{T}$ strains, two sequences annotated as potential RpoD transcription factors were found

**References**

Amarelle V, Roldán DM, Fabiano E, Guazzaroni M-E (2023) Synthetic Biology Toolbox for Antarctic Pseudomonas sp. Strains: Toward a Psychrophilic Nonmodel Chassis for Function-Driven Metagenomics. ACS Synth Biol. https://doi.org/10.1021/acssynbio.2c00543

Bagdasarian M, Lurz R, Rückert B, et al (1981) Specific-purpose plasmid cloning vectors II. Broad host range, high copy number, RSF 1010-derived vectors, and a host-vector system for gene cloning in Pseudomonas. Gene 16:237–247. https://doi.org/10.1016/0378-1119(81)90080-9

Crooks GE, Hon G, Chandonia J-M, Brenner SE (2004) WebLogo: A Sequence Logo Generator. Genome Res 14:1188–1190. https://doi.org/10.1101/gr.849004

Ditta G, Stanfield S, Corbin D, Helinski DR (1980) Broad host range DNA cloning system for gram-negative bacteria: construction of a gene bank of Rhizobium meliloti. Proc Natl Acad Sci 77:7347–7351. https://doi.org/10.1073/pnas.77.12.7347

Ferrés I, Amarelle V, Noya F, Fabiano E (2015) Identification of Antarctic culturable bacteria able to produce diverse enzymes of potential biotechnological interest. Adv Polar Sci 26:71–79. https://doi.org/10.13679/j.advps.2015.1.00071

Hall A, Donohue T, Peters J (2023) Complete sequences of conjugal helper plasmids pRK2013 and pEVS104. microPublication Biol 2023:. https://doi.org/10.17912/micropub.biology.000882

Hanahan D (1983) Studies on transformation of Escherichia coli with plasmids. J Mol Biol 166:557–580. https://doi.org/10.1016/S0022-2836(83)80284-8

Meade HM, Long SR, Ruvkun GB, et al (1982) Physical and genetic characterization of symbiotic and auxotrophic mutants of Rhizobium meliloti induced by transposon

Tn5 mutagenesis. J Bacteriol 149:114–122. https://doi.org/10.1128/jb.149.1.114-122.1982

Sawana A, Adeolu M, Gupta RS (2014) Molecular signatures and phylogenomic analysis of the genus Burkholderia: proposal for division of this genus into the emended genus Burkholderia containing pathogenic organisms and a new genus Paraburkholderia gen. nov. harboring environmental species. Front Genet 5:. https://doi.org/10.3389/fgene.2014.00429

Silva-Rocha R, Martínez-García E, Calles B, et al (2013) The Standard European Vector Architecture (SEVA): a coherent platform for the analysis and deployment of complex prokaryotic phenotypes. Nucleic Acids Res 41:D666–D675. https://doi.org/10.1093/nar/gks1119

Vandamme P (2004) Taxonomy of the genus Cupriavidus: a tale of lost and found. Int J Syst Evol Microbiol 54:2285–2289. https://doi.org/10.1099/ijs.0.63247-0
